# Prediction of cellular morphology change under perturbations with transcriptome-guided diffusion model

**DOI:** 10.1101/2025.07.16.665106

**Authors:** Xuesong Wang, Yimin Fan, Yucheng Guo, Chenghao Fu, Kinhei Lee, Khachatur Dallakyan, Yaxuan Li, Qijin Yin, Yu Li, Le Song

## Abstract

Investigating the cell morphology change after perturbations with high-throughput image-based profiling is of growing interest, considering its wide applications in phenotypic drug discovery, including MOA (Mechanism Of Action) prediction, compound bioactivity prediction, and drug repurposing. However, the vast space of chemical and genetic perturbations makes it infeasible to fully explore all the potential perturbations with image-profiling technologies. Consequently, developing a powerful in-silico method to simulate high-fidelity cell morphological response under perturbations can reduce the experiment costs and accelerate drug discovery. Motivated by this, we proposed MorphDiff, a transcriptome-guided latent diffusion model for accurately predicting the cell morphology response to perturbations. We applied MorphDiff to two large-scale datasets, including one drug perturbation and one genetic perturbation cell morphology dataset covering thousands of diverse perturbations. Extensive benchmarking and comparison with baseline methods show the remarkable accuracy and fidelity of MorphDiff in predicting cell morphological changes under unseen perturbations. Furthermore, we explored the utilities of MorphDiff in identifying and retrieving the MOAs of drugs, which is a crucial application in phenotypic drug discovery. With the designed pipeline for MOA retrieval, we demonstrated MorphDiff’s capability to boost the retrieval of the drugs’ MOAs (Mechanism Of Actions) by generating realistic cell morphology profiles. The average MOA retrieval accuracy of MorphDiff-generated morphology is comparable with that of the ground truth cell morphology, and consistently outperforms the baseline method and gene expression-based retrieval by 29.1% and 9.7% respectively. We also validated that complementary information provided by cell morphology generated by MorphDiff can help discover drugs with dissimilar structures but the same MOAs. In summary, with its strong capabilities in generating high-fidelity cell morphology on unseen perturbations, we envision MorphDiff as a powerful tool in phenotypic drug discovery by accelerating the phenotypic screening of vast perturbation space and improving MOA identification.

## 1 Introduction

Characterizing and predicting the cell states under genetic and drug perturbations remains one of the most challenging and meaningful directions in the single-cell biology domain. By understanding how individual cells respond to specific genetic and drug perturbations, researchers can better map the pathways that contribute to particular cell states and identify the potential targets for therapeutic intervention. Multiple aspects of cell states may change under perturbations, including but not limited to gene expression profile, proteomic composition, metabolic status, and cell morphology. Out of various modalities of cell states, the investigation of cell morphology changes following perturbations through high-throughput image-based profiling has gained significant interest due to its broad applications in phenotypic drug discovery, including the prediction of Mechanism of Action (MOA), compound bioactivity, and drug repurposing [1].

However, considering the vast number of synthesizable chemical compounds and genes in perturbation space, it is infeasible to profile the cell morphology for all possible perturbations. Therefore, developing an in-silico method for simulating the cell morphological response under perturbations emerges as a challenging and meaningful topic [2]. With a powerful and reliable tool to infer the cell morphology change under perturbations, exploring the vast perturbation space can be significantly accelerated, further promoting downstream drug discovery pipelines, such as MOA prediction. Computational approaches have been proposed to address this challenge [3]. However, the accuracy and fidelity of these methods remain inadequate. First, as indicated by existing work IMPA [3], prediction on unseen perturbations will only perform well if the model has seen a similar perturbation with a similar effect in the training dataset. As structurally similar drugs or co-expression gene knockout may have distinct effects [4], how to faithfully encode and represent perturbations is critical to improving the prediction generalizability on unseen perturbations. Second, besides perturbations, cellular morphology is influenced by a wide range of factors, including but not limited to batch and well position effects [5]. Thus, the morphology data can be pretty noisy.

Motivated by this, we developed MorphDiff, a scalable transcriptome-guided diffusion model for predicting the cell morphology change to unseen perturbations. MorphDiff predicts the perturbed cell morphology with the perturbed L1000 gene expression profile as the condition. The following motivations inspire the choice of using gene expression profiles as the condition. Firstly, gene expression plays a crucial role in determining cell morphology by directing the synthesis of proteins that regulate cellular structure and dynamics [6; 7]. Though the relationship between cell morphology and gene expression is complex, there is still shared and complementary information between them [8]. Therefore, the perturbed gene expression is more informative in determining cell morphology than prior knowledge-guided encoding of perturbation such as drug SMILES representation and Gene2vec [9] embedding. Secondly, the L1000 assay offers a much larger pool of publicly available datasets compared with cell morphology profiling [10]. Thus, obtaining the gene expression of cells treated with unseen perturbations is much easier than obtaining the cell morphology, enabling our tool to be applicable in broader scenarios.

The architecture of MorphDiff is based on the Latent Diffusion Model [11], an advanced generative model inspired by thermodynamics that learns to reverse the data from an unordered state to an ordered state. We employ a Variational Autoencoder (VAE) [12] to compress cell morphology into low-dimensional embeddings and subsequently train a latent diffusion model [11] using gene expression profiles as the condition. Compared with traditional generative models such as Generative Adversarial Network (GAN) [13] used in MorphNet [14] and IMPA [3], the diffusion-based architecture has the following advantages. Firstly, the latent diffusion model is highly robust to noise as, by design, diffusion models transform data into a noisy state and learn to reconstruct it, which makes it highly suitable for cell morphology datasets. Secondly, the latent diffusion model supports flexible conditions, further extending the potential utility of MorphDiff. Besides solely relying on the perturbed transcriptome as input, the pre-trained MorphDiff model can also take an unperturbed cell morphology as input with the perturbed cell transcriptome as the condition to infer the continuous transition of cell morphology from unperturbed to perturbed without additional training. Thirdly, diffusion models are more advanced on normal image synthesis tasks than other GAN-based generative models [15].

We conducted extensive experiments to demonstrate the utility of MorphDiff in facilitating the exploration of cell morphological response towards unseen perturbations. First, on two curated large-scale datasets encompassing diverse genetic perturbations and drug perturbations, MorphDiff achieves state-of-the-art performance in standard image generation assessment metrics compared with a wide range of baseline methods [14; 3; 16; 17; 18]. Furthermore, we evaluated the generated output of MorphDiff by incorporating CellProfiler [19] and DeepProfiler [20], which can be used to extract morphological features and embeddings. Our findings suggest that the interpretable morphological features extracted from the generated output of MorphDiff are much closer to the features extracted from ground truth perturbed morphology, which further illustrates the high fidelity of MorphDiff in predicting cell morphological response to unseen perturbations. Meanwhile, MorphDiff well captures the intricate correlation between transcriptome and morphology change under perturbations. In terms of the correlation between transcriptome-generated cell morphology association and ground truth transcriptome-morphology association, MorphDiff outperforms the baseline method by 75%.

To further explore potential applications of MorphDiff in phenotypic drug discovery, we focused on an essential direction in drug discovery: MOA retrieval of drugs. Previous approaches to elucidate drug MOA include drug structure analysis, literature mining, and transcriptome response analysis [6]. We revealed that cell morphology provides crucial information in identifying drug MOA, and more importantly, the cell morphology inferred by MorphDiff on unseen drug perturbations can also benefit MOA identification. With the designed MOA retrieval pipeline, we found that the generated cell morphology from MorphDiff achieves an excellent performance comparable with ground truth cell morphology, and consistently, the average accuracy of Morphdiff-generated output surpasses the baseline method and gene expression-based retrieval by 29.1% and 9.7% respectively. More importantly, we validated that the complementary information provided by cell morphology generated by MorphDiff can help discover drugs with dissimilar structures but similar MOAs, such as *Diclofenamide* and *fraxidin methyl ether*, which have distinct structures but both annotated with *Carbonic anhydrase inhibitor* MOA. In summary, MorphDiff is a powerful tool in phenotypic discovery by accurately generating cell morphology under unseen perturbations, with promising applications in facilitating the exploration of vast phenotypic perturbation screening space and assisting in determining the MOAs of structurally diverse drugs.

## 2 Results

### 2.1 Method Overview

MorphDiff maps L1000 gene expression to cell morphology images through a Latent Diffusion Framework [11]. Concretely, as shown in Figure 1 a, paired L1000 gene expression and cell morphology images are curated for the same perturbations. The cell morphology images are acquired by the Cell Painting assay and are usually composed of five channels, including DNA, ER, RNA, AGP, and Mito. MorphDiff is made up of two main components (Figure 1 b), Morphology VAE (MVAE) and Latent Diffusion Model (LDM). MVAE consists of the encoder and decoder parts. The encoder takes the cell morphology images composed of five channels as input and outputs the latent representation, while the decoder reconstructs the original input image based on the latent representation. The Latent Diffusion Model (LDM) consists of the noising process and denoising process. In the noising process, Gaussian random noise is sequentially added to the latent representation of morphology from 0 to *T*. The final *Z*_*T*_ follows the standard Gaussian distribution. In the denoising process, LDM is trained to recursively remove the noise from *Z*_*t*_ conditioned on L1000 gene expression as *t* decreases from *T* to 0. LDM is implemented with denoising U-Net architecture augmented with attention mechanism [22]. As a condition, L1000 gene expression is combined with the key and value of the attention mechanism in LDM. In practice, the training of LDM is equivalent to minimizing the variational upper bound. Details are described in Section 4.2. As illustrated in Figure 1 c, pre-trained MorphDiff can be applied in two ways. In the first mode, MorphDiff takes the L1000 gene expression as the condition and denoises the corresponding cell morphology images from random noise distribution. In the second mode, MorphDiff takes the L1000 gene expression for one specific perturbation as the condition and transforms the morphology images from the control set to the predicted morphology images for that perturbation. Compared with previous works in generating cell morphology images, MorphDiff is the only tool that supports generation from gene expression to morphology and transformation from unperturbed morphology to perturbed morphology. Meanwhile, our framework corporates CellProfiler [19] and DeepProfiler [20] to generate biological meaningful CellProfiler morphological features and DeepProfiler morphological embeddings, thereby exhibiting better interpretability. Among all the methods in Figure 1 d, only MorphDiff and IMPA are able to generate five-channel images, which can be used to extract morphological features and morphological embeddings by CellProfiler and DeepProfiler. Therefore, the analysis and comparison involving morphological features and morphological embeddings in the following article only involve these two methods. We collected two largescale cell morphology image datasets for the model training evaluation spanning 960 drug perturbations and 130 genetic perturbations. We split the dataset into three parts: training set, in-distribution (ID) set, and out-of-distribution (OOD) set. We selected 10% perturbations as the OOD set (96 drug perturbations and 13 genetic perturbations). For the remaining perturbations, we performed a 90%:10% split for the cell morphology images for each perturbation with 90% as the training set and 10% as the ID set. Therefore, the training and ID set covers the same perturbations but with different cell morphology images. For benchmarking and downstream task, we generated gene embeddings and drug embeddings when implementing baseline methods following the requirements and conducting the analysis of drug similarity. The gene embeddings are generated with [9] and drug embeddings are generated with RDKit [23].

**Figure 1.**
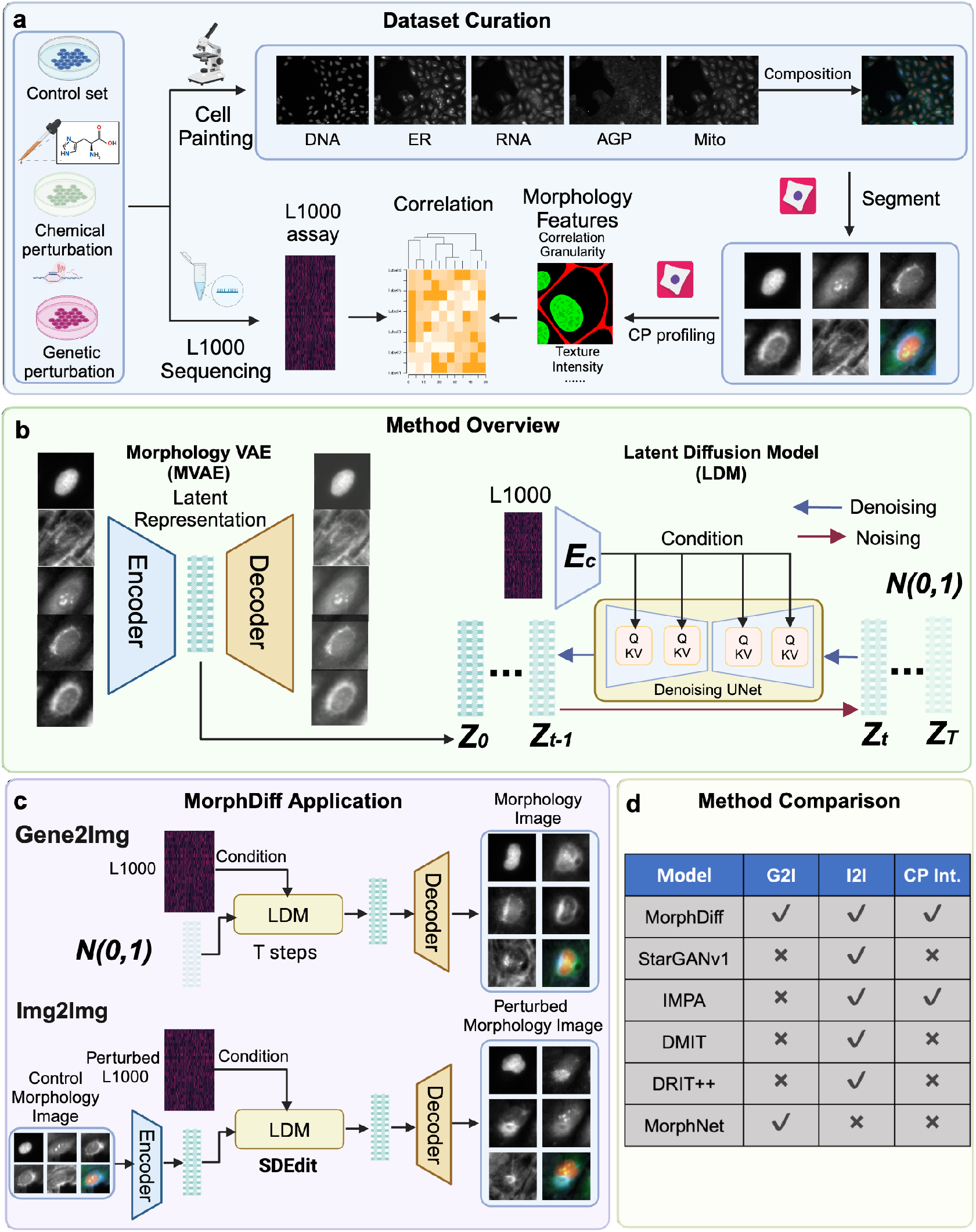
Overview of the MorphDiff Framework. **a.** In this study, the multimodal dataset of U2OS cells consists of three parts: control set (DMSO), genetic perturbation set (130 unique perturbations), and compound treatment set (960 unique perturbations). For each unique perturbation, the morphology images with five channels are collected using CP (Cell Painting), and the gene expressions are collected using L1000 assays. CellProfiler is then used to segment the cells and extract CellProfiler features at the single-cell level. Morphology images and gene expression together characterize the response of U2OS cells to specific perturbations. **b.** MorphDiff is composed of two main components: Morphology VAE (MVAE) and Latent Diffusion Model (LDM). The MVAE encoder encodes the multi-channel cell morphology images into latent representation, and the decoder reconstructs the original cell morphology images based on the latent representation. LDM is trained to denoise from Gaussian random noise *Z*_*T*_ to morphology latent representation *Z*_0_ recursively conditioned on L1000 gene expression. **c.** MorphDiff can be applied in two ways to generate cell morphology images with perturbations: L1000 gene expression to cell morphology generation (G2I) and perturbed L1000 gene expression combined with control morphology images to perturbed cell morphology images generation (I2I). I2I is implemented with SDEdit [21] without re-training. **d.** Comparison of MorphDiff with other related tools. Created with BioRender.com.

### 2.2 MorphDiff accurately predicts changes in cell morphology with genetic perturbations

To illustrate the superiority of MorphDiff, we benchmarked the general prediction performance on the OOD set of genetic perturbations data. We compared MorphDiff with a wide range of baseline methods, including MorphNet [14], DMIT [16], DRIT++ [17], StarGANv1 [18] and IMPA [3]. Some baseline methods, such as IMPA, DRIT++, DMIT, and StarGANv1, do not utilize the gene expression and model the perturbation process as an image-to-image translation problem. At the same time, MorphNet performs direct gene expression to morphology image translation. To ensure the statistical reliability of the results, we sampled 10 times for each method with different random seeds and conducted unpaired Welch t-tests for each comparison. It should be noted that one star (‘*’) means MorphDiff surpasses the baseline method with a p-value less than 0.05, while two stars and three stars correspond to p-values less than 0.01 and 0.001, respectively. On the contrary, hashtag (‘#’) represents the situation where the baseline method outperforms MorphDiff. The corresponding relationship between the p-value and the number of hashtags is the same as the star. As depicted in Figure 2 a MorphDiff consistently outperforms all other methods on the fid metric, which is the most prevalent and significant metric for evaluating generative models. In terms of other metrics, MorphDiff generally ranks the best or the second best and achieves the highest average score.

**Figure 2.**
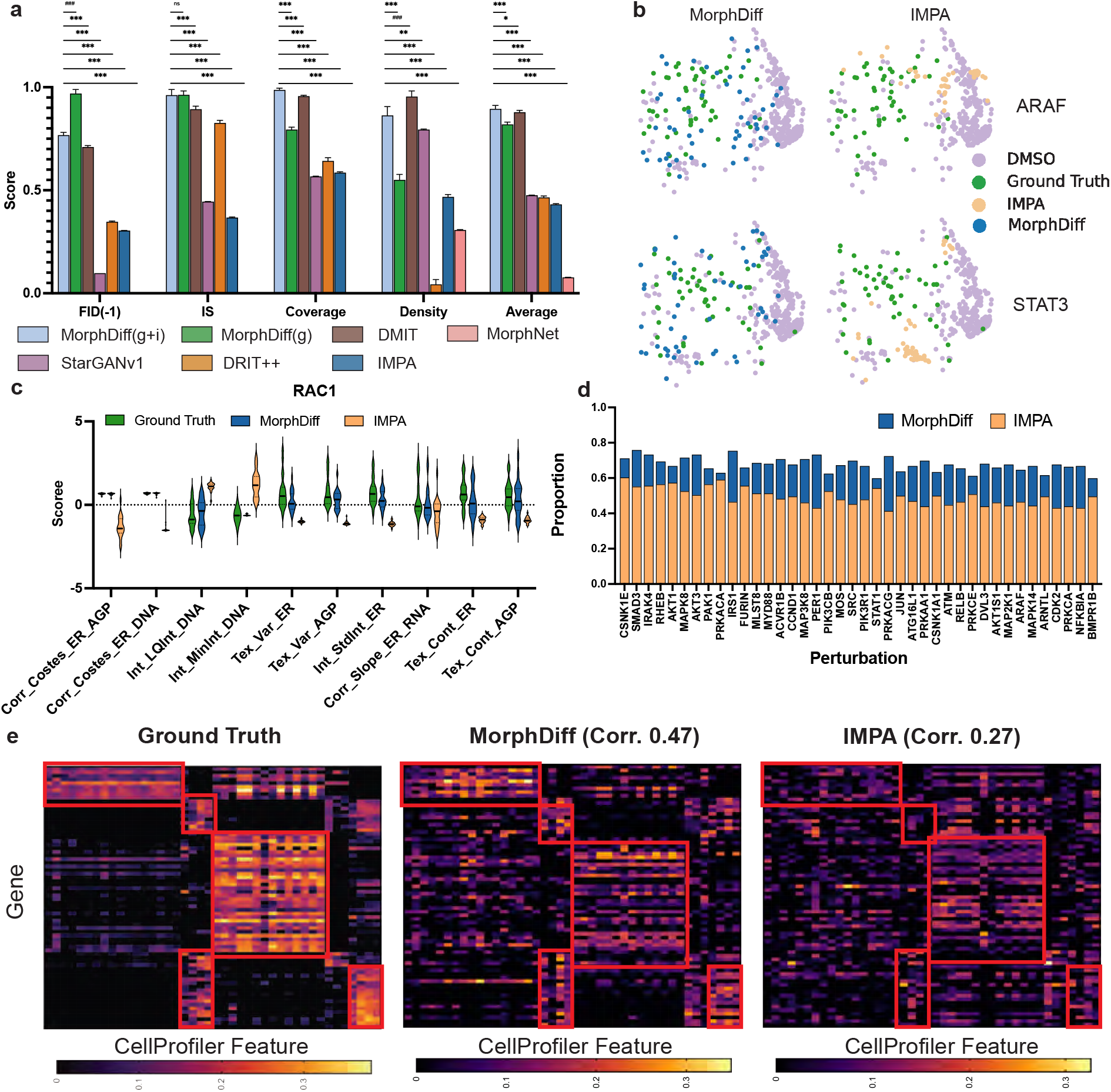
MorphDiff predicts changes in cell morphology with genetic perturbations. **a.** Performance benchmarking on unseen genetic perturbations. MorphDiff(g) denotes generating the perturbation morphology images with only the perturbation gene expression. MorphDiff(g+i) denotes generating the perturbation morphology with the perturbation gene expression and a control reference image. Linear normalization was conducted on the reciprocal of fid and the other three metrics separately to convert their values to the range of 0 to 1. A larger value indicates better performance. **b.** UMAP visualization of the generated cell morphology CellProfiler feature on two perturbations for IMPA and MorphDiff. **c.** Distribution of CellProfiler features between ground truth, generated morphology from IMPA, and generated morphology from MorphDiff on RAC1 genetic perturbation. E stands for ERSyto. EB stands for ERSytoBleed. H stands for Hoechst. Pg stands for Ph golgi. LQ stands for LowerQuartile. Var stands for Variance. Tex stands for Texture. Corr stands for correlation. Int stands for Intensity. **d.** The t-test results to test the difference between the generated cell morphology CellProfiler features and the ground truth CellProfiler features. The y-axis indicates the proportion of generated CellProfiler features not significantly differing from ground truth CellProfiler features through an independent t-test (*p <* 0.05). Across different types of perturbations, morphology generated by MorphDiff consistently has more CellProfiler features that do not have significant differences with ground truth, which indicates the high generation quality of MorphDiff. **e.** The heatmap of the correlation between the CellProfiler features of ground truth morphology, MorphDiff-generated morphology, and IMPA-generated morphology with L1000 gene expression.

In addition, we visualized the generated output of MorphDiff along with other baseline methods to intuitively demonstrate the superiority of our method. We used the results of MorphDiff (G2I) generation output. In Appendix Figure 1, we visualized the generated samples on four genetic over-expression perturbations (AKT1, AXIN2, JUN, STK11) and the ground truth samples. We can observe that the generated samples of MorphNet are of relatively low quality and can not be distinguished across perturbations. The images generated by DRIT++ exhibit high variance across different perturbations, but the quality of the generated images is also low. StarGAN, DMIT and IMPA can generate relatively high-quality images, but the generated morphologies are unsatisfactory on some perturbations such as AXIN2 and JUN. The visual quality of the images generated by the proposed method MorphDiff is the highest, and the generated images are the most visually similar to the ground truth images.

Furthermore, we performed an in-depth analysis of whether MorphDiff can accurately predict changes in cell morphology across a wide range of genetic perturbations on the ID set of genetic perturbations data. We performed quantitative analysis using the CellProfiler features to demonstrate the effectiveness of MorphDiff. CellProfiler features are a set of morphological features extracted with CellProfiler related to cell morphology, measuring the texture, intensity, and granularity inside each morphology channel and the correlation across different morphology channels. The generation setting of MorphDiff in this analysis is *Gene2Img*. In Figure 2 b, we used UMAP to project the morphological features into two dimensions for visualization. We selected the most important 10 CellProfiler features discriminative of the perturbation types to form the morphological feature vector. The details of the feature selection strategy are discussed in Section 4.3 (Morphological feature selection based on discriminability of perturbations). We can observe that the distribution of ground truth perturbated cell morphology also has an apparent variance considering the variance in individual cell development. The generated outputs of IMPA form a dense cluster and are far away from the ground truth distribution. In contrast, the outputs of MorphDiff are much more diverse and resemble the ground truth distribution. We also conducted the same analysis on other genes, and the results in Appendix Figure 2 display a consistent effect. Then, we turned to a more fine-grained analysis of how well the generated morphology aligns with the ground truth morphology regarding the values of individual CellProfiler features. In Figure 2 c, we visualized the values of the 10 most important cell morphology features of the ground truth perturbation morphology, morphology generated by MorphDiff, and morphology generated by IMPA on RAC1 over-expression. We can see that MorphDiff-generated morphology aligns well with the ground truth morphology compared with the IMPA-generated morphology. Results on more perturbations are shown in Appendix Figure 3 and Appendix Figure 4. To have a more holistic view of how close the MorphDiff-generated morphology aligns with the ground truth distribution, we performed statistical testing for a more quantitative comparison. Concretely, for each feature out of all CellProfiler features extracted, we conducted an independent t-test across the ground truth perturbation CellProfiler features and the generated CellProfiler features. Suppose the p-value of the t-test is larger than 0.05. In that case, we conclude that insufficient evidence shows significant differences between the generated perturbation morphological features and the ground truth perturbation morphological features. As shown in Figure 2 d, the proportion of generated CellProfiler features that align with the ground truth features of MorphDiff is much larger than IMPA. For MorphDiff, more than seventy percent of generated CellProfiler morphology features align with the ground truth perturbation CellProfiler features, which demonstrates the effectiveness of MorphDiff in predicting accurate perturbation morphology.

**Figure 3.**
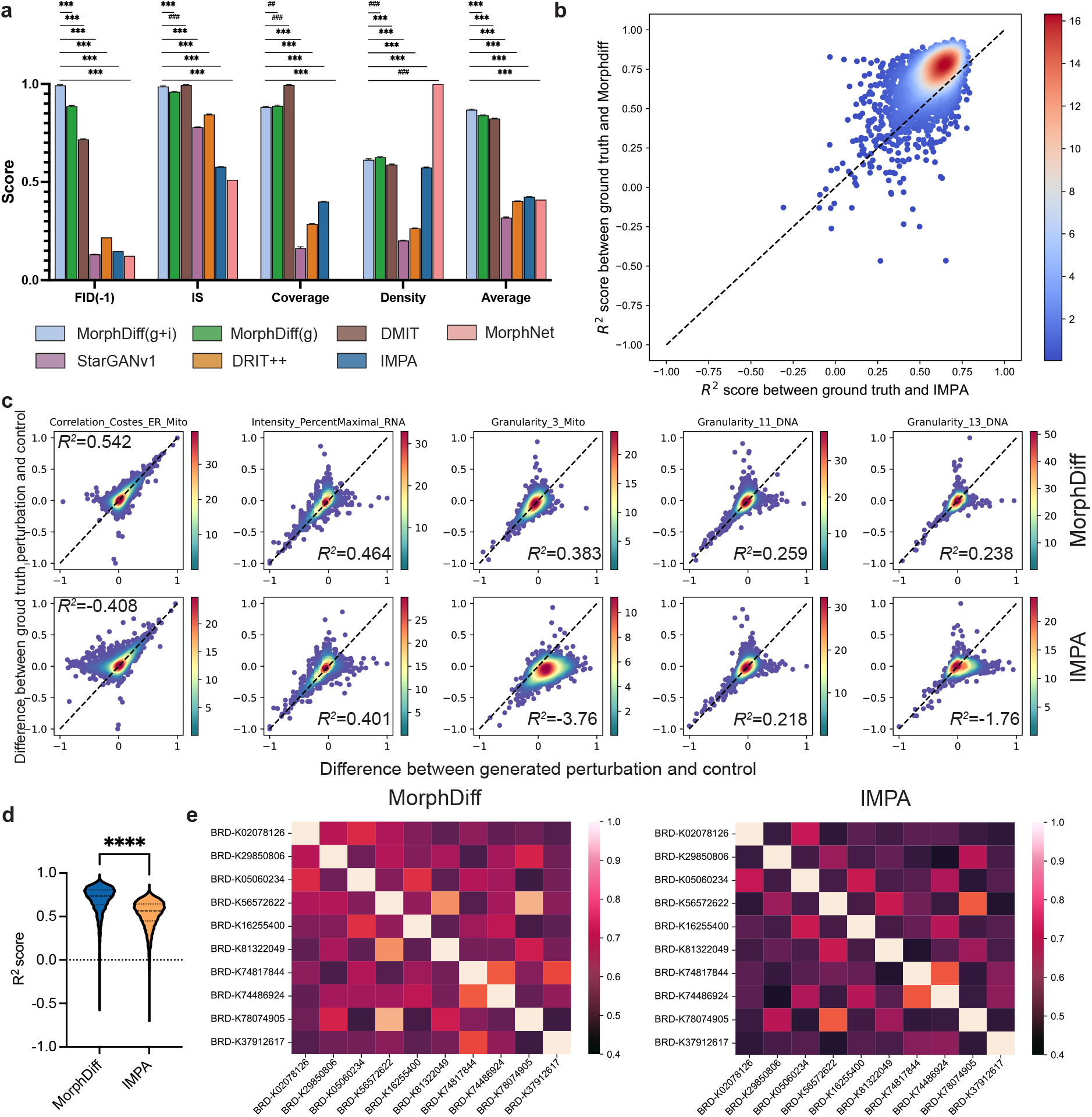
MorphDiff predicts changes in cell morphology after drug perturbations. **a.** General generative performance on Drug OOD set. Linear normalization was conducted on the reciprocal of fid and the other three metrics separately to convert their values to the range of 0 to 1. Each method was sampled 10 times with different seeds, and an unpaired Welch t-test was conducted to ensure the significance of the results. **b.** The *R*^2^ score between the ground truth CellProfiler feature vectors and the generated CellProfiler feature vectors on the Drug ID set. Most points are concentrated above the diagonal, indicating that the morphology generated by MorphDiff closely resembles the ground truth. **c.** The difference between control and perturbation for selected CellProfiler features on the Drug ID set. The x-axis displays the difference between the generated perturbed and the control CellProfiler features, while the y-axis displays the difference between the ground truth perturbed and the control CellProfiler features. The higher the point density along the diagonal, the closer the generated difference is to the ground truth. More results can be found in the Appendix Figure 8. The score represents *R*^2^ score measuring the similarity between the predicted difference and ground truth difference. **d.** The *R*^2^ scores, measuring the similarity of CellProfiler features between the ground truth and the generated samples on the Drug OOD set. **e.** The relative accuracy comparison between MorphDiff and IMPA reveals that MorphDiff outperforms IMPA in almost all pairs of OOD perturbations.

**Figure 4.**
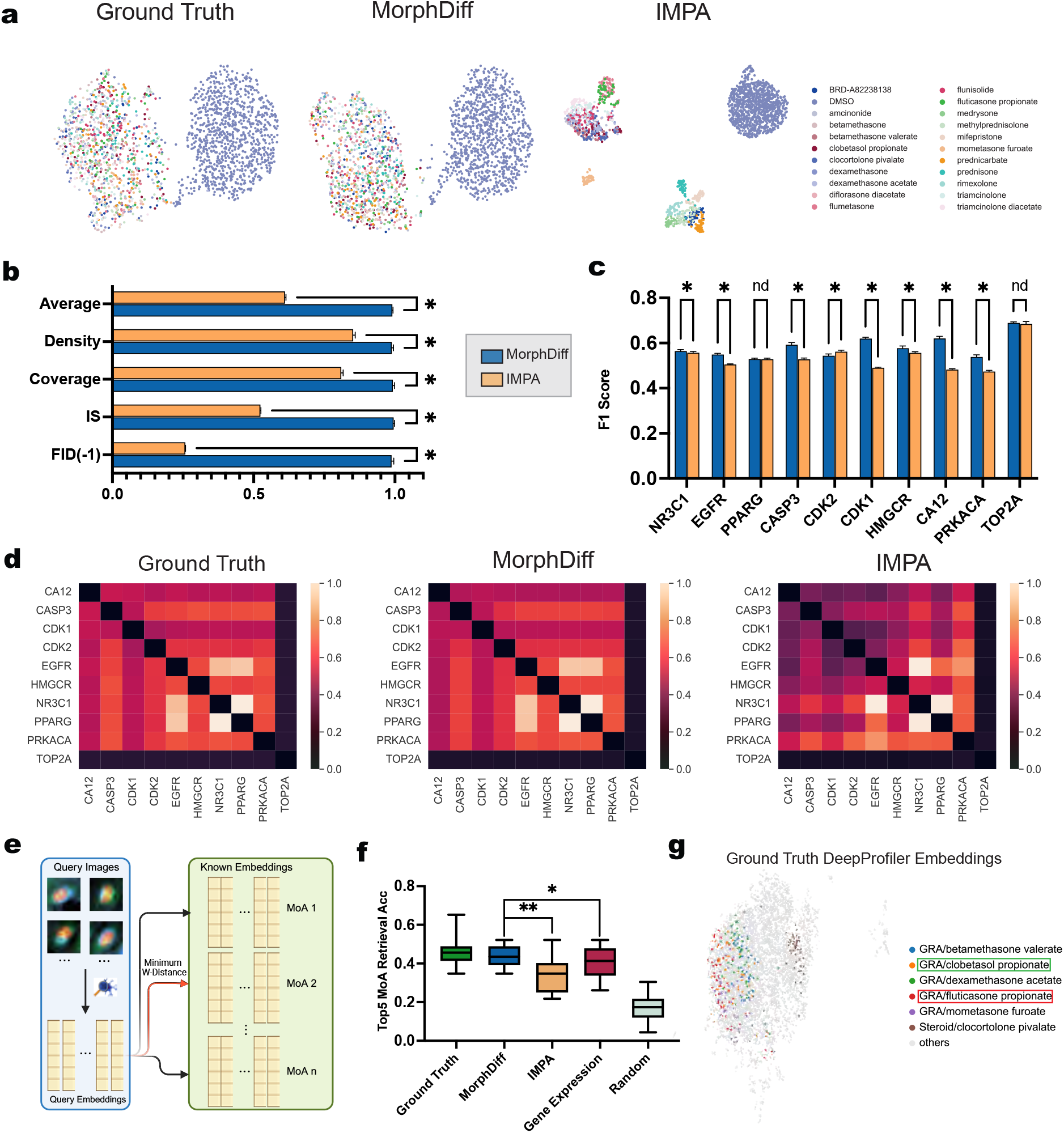
Pre-trained MorphDiff potentially promotes drug discovery. **a.** CellProfiler feature vectors extracted from both ground truth and generated images. The blue points represent control CellProfiler feature vectors, while other colors represent CellProfiler feature vectors perturbed by drugs targeting the *NR3C1* gene. **b.** General generative evaluation of MorphDiff and IMPA. **c.** F1 score measures the performance of two methods in identifying whether more than 200 CellProfiler features will undergo significant changes under drug perturbations for various targets. **d.** W-distance between various pairwise target-level perturbations for ground truth and generated images, computed with DeepProfiler embeddings extracted via DeepProfiler [20]. Upon normalization, the results indicate that MorphDiff aligns consistently with the ground truth, whereas IMPA does not. **e.** Schematic overview of retrieval workflow. **f.** Top 5 Mechanisms of Action (MOA) retrieval results for different modalities. Ground truth denotes retrieval with ground truth morphological embeddings, while MorphDiff and IMPA signify retrieval with respectively generated DeepProfiler embeddings. Gene Expression represents retrieval according to the MSE (Mean Square Error) between gene expression. **g.** Ground Truth DeepProfiler embeddings projected with UMAP. The drug in the green box is *clobetasol propionate* annotated with GRA, and the drug in the red box are the closed drugs based on Wasserstein distance in morphological embedding space. GRA and Steroid are two types of MOAs. GRA is the abbreviation of *Glucocorticoid Receptor Agonist*.

We also validated the effectiveness of MorphDiff in capturing the correlation between gene expression and morphology. For better clarity, we selected CellProfiler features that MorphDiff and IMPA can best recover. The details of feature selection are described in Section 4.3. Figure 2 e shows the correlation between gene expression and CellProfiler features of ground truth, MorphDiff, and IMPA, respectively. In Figure 2 e, we observed that several groups of genes and CellProfiler features have a high correlation with each other. For example, the group on the top left of the heatmap mainly consists of features related to Correlation and Intensity associated with ER, Mito, and AGP channel (Appendix Figure 5). Another group on the bottom right mainly consists of features related to the intensity of DNA and RNA channels (Appendix Figure 5). The details correlation of MorphDiff and IMPA can be found in Appendix Figure 6 and Appendix Figure 7. We computed the correlation between the correlation map of the generated samples and ground truth samples. We observed that MorphDiff-generated samples have a much higher correlation (0.47) than IMPA-generated samples (0.27). Therefore, MorphDiff captures the pattern of cell morphological feature changes across different genes well, while IMPA fails to capture the correlation pattern between gene expression and cell CellProfiler features.

### 2.3 MorphDiff captures morphological changes of cell morphology across thousands of drugs treatment

We first benchmarked all methods on the Drug OOD set with general evaluation metrics, in which the meanings of stars and hashtags are the same as in Section 2.2. As Figure 3 a shows, MorphDiff ranks first on the average score. It is worth noting that MorphNet achieves a remarkably high density score but nearly zero coverage score. Upon checking the images generated by MorphNet, we found that it produced almost identical images as the part of training images. As there exist some drugs in the OOD set which may yield similar effects as those in the training dataset, the generated images may always lie in the neighbors of real images treated with similar drugs as the training set, which results in a high density score as described in the definition of density score in Section 4.5. However, as MorphNet produces images that are highly similar to those of the training dataset, it fails to make reliable predictions on those drugs with distinct effects compared with the training set, which leads to a meager coverage score. This is a common problem of GAN-based model, as known as mode collapse [24].

To investigate the response of cell morphology to small molecular compounds from a more fine-grained and interpretable perspective, we extracted the CellProfiler features from the generative morphologies and compared them with features extracted from ground truth morphologies. Concretely, we extracted CellProfiler features from 3000 ground truth images sampled from the Drug ID set and corresponding images generated by MorphDiff (I2I) and IMPA. Over 200 informative CellProfiler features are extracted and analyzed. We calculated *R*^2^ scores between the ground truth CellProfiler feature vectors and the corresponding generated CellProfiler feature vectors. A higher *R*^2^ score indicates better proximity between the generated and the ground truth CellProfiler feature vectors. We compared the *R*^2^ score of MorphDiff versus ground truth (y-axis) and IMPA versus ground truth (x-axis) in Figure 3 b, with each point indicating a unique cell morphology. Figure 3 b shows that most scatter points lie above the diagonal, signifying that MorphDiff outperforms IMPA in most of the 3000 samples. Meanwhile, the y-coordinate corresponding to the highest density is close to one. This suggests that MorphDiff can generate highly realistic samples using a real control image as a base image.

Furthermore, we examined the effectiveness of our tool in predicting the features that undergo the most significant changes after perturbation. For each CellProfiler feature, we conducted a chi-square test between ground truth control CellProfiler features and ground truth perturbed CellProfiler features to determine whether this feature will significantly change after perturbation. We picked the features with p-values less than 0.05 as significantly changed and calculated the *R*^2^ score between the real and generated samples for each feature across 3000 images. After that, we found that MorphDiff outperforms IMPA on 69 out of 101 significantly changed features. To intuitively demonstrate the superiority of MorphDiff, we displayed parts of features by using a scatter plot to illustrate the degree of proximity between ground truth and generated images as shown in Figure 3 c and Appendix Figure 8 a. The x-axis represents the difference between generated CellProfiler features and corresponding control CellProfiler features, while the y-axis represents the difference between ground truth and control CellProfiler features. Considering the high density on the diagonal, it is evident that MorphDiff can accurately predict morphological changes on parts of significantly changed CellProfiler features.

To further evaluate the generalizability of MorphDiff, we performed the same analysis on the Drug OOD set. We selected 10 OOD perturbations with the most images, resulting in 4828 images. On the individual sample level, Figure 3 d demonstrates that most samples generated by MorphDiff are more similar to the ground truth compared to samples generated by IMPA on the OOD set, and *R*^2^ scores of MorphDiff-generated CellProfiler feature vectors are significantly higher than baseline method based on unpaired Welch t-test with p-value less than 0.0001. For CellProfiler features, consistent with the results on the ID set, MorphDiff outperforms IMPA on 101 out of 128 significantly changed CellProfiler features and performs well on parts of the significantly changed CellProfiler features, as shown in Appendix Figure 8 b. As we release the code used to generate these figures, people can check the *R*^2^ of all significantly changed CellProfiler features with our Jupyter file.

Moreover, assessing whether the model can capture the diversity between different perturbations and generate perturbed cell morphology with high perturbation specificity is essential. We utilized DeepProfiler [20] to extract morphological embeddings from each image, and we term them as DeepProfiler embeddings. Subsequently, we trained an SVM binary classifier[25] for each pair of perturbations and calculated the classification accuracy for DeepProfiler embeddings generated by MorphDiff, IMPA, and the ground truth images. We computed the relative accuracy by dividing the accuracy obtained from the ground truth when comparing the performance of MorphDiff and IMPA. The results in Figure 3 e clearly show that MorphDiff achieves higher relative accuracy than IMPA. MorphDiff achieves a relative accuracy of more than 0.7 in over half of the situations, with the highest relative accuracy around 0.9. In contrast, IMPA consistently achieves lower accuracy, with most cases falling below 0.7 and many cases below 0.5. The high classification accuracy between multiple perturbations implies the superiority of MorphDiff in capturing the subtle diversity between different perturbations.

### 2.4 Pre-trained MorphDiff as a promising tool in phenotypic drug discovery

Small molecule compounds are agents that modify a target to affect its functionality [26]. These compounds can either reduce or accelerate the activity of a target, which is typically a biomolecule like a protein, enzyme, receptor, or gene. These biomolecules are involved in signaling or metabolic pathways, often specific to diseases [27]. Biomolecules play a pivotal role in disease development and progression, primarily through communication facilitated by interactions between proteins and nucleic acids or proteins themselves. These interactions often lead to signal amplification or metabolic process alterations, further affecting disease development. Typically, a target can be associated with multiple drugs. For instance, *amcinonide, dexamethasone*, and *betamethasone* can all impact the gene *NR3C1*, while *LFM-A12, RG-14620, WHI-P154*, and *chrysophanol* can affect the gene *EGFR*. Based on these observations, we aim to investigate whether drugs acting on the same target will exhibit similar influences on morphological changes, and we wonder whether MorphDiff can capture such patterns. We selected 10 targets (Section 4.3 Target dataset) with sufficient corresponding compounds to include enough cell morphology for analysis and these data are not present in the training set for MorphDiff and IMPA. To determine whether the images of perturbations related to these targets are significantly different from control unperturbed images, we extracted CellProfiler feature vectors from real images and projected the CellProfiler features through UMAP [28], as depicted in Figure 4 a, using the target *NR3C1* as an example. We observed that the ground truth CellProfiler feature vectors induced by 21 drugs are clearly distinct from the real control set in the CellProfiler feature space. Moreover, these CellProfiler feature vectors cluster together, indicating that these drugs acting on the same target cause a similar effect in the CellProfiler feature space, which we term as target-level effect. To explore whether MorphDiff can capture such target-level effect, we used the control images and gene expression perturbed by these 21 types of drug as input and then applied the pre-trained MorphDiff (pre-trained with the training set of drug perturbation data, which does not intersect with Target dataset) to generate corresponding morphology images. As shown in Figure 4 a, the CellProfiler feature vectors generated by the pre-trained MorphDiff drive a consistent effect with ground truth after UMAP projection. On the other hand, IMPA generates multiple clusters of perturbations targeting the same gene, while these perturbations should have a similar effect according to the ground truth, which indicates its poor generalization performance. We did the same analysis on the other 9 targets and the results can be found in Appendix Figure 9 and Appendix Figure 10. Additionally, drugs that act on the same targets may have contrasting effects, potentially causing different impacts on cell morphology due to different Mechanisms of Action (MOAs), such as inhibitors and agonists. However, from the UMAP visualization of the ground truth cell morphology, the diversity is not as obvious as expected.

To quantitatively assess the generalization capabilities of MorphDiff, we conducted a benchmark test on the pre-trained IMPA and MorphDiff using the Target dataset. We randomly sampled 10 times with different random seeds. As illustrated in Figure 4 b, MorphDiff significantly surpasses IMPA with the p-value less than 0.05 based on an unpaired Welch t-test for each image generation quality metric. This result underscores the exceptional generalization ability of MorphDiff. Moreover, we investigated whether these CellProfiler features undergo significant changes when exposed to drugs that target different genes. Therefore, we adopted the chi-square test on the same CellProfiler features between control and perturbed morphology to identify significantly affected CellProfiler features with the p-value threshold at 0.05. With the significantly affected CellProfiler features identified in ground truth morphology as ground truth label, we calculated the F1 score (details in Section 4.5) for CellProfiler features identified in morphology generated by pre-trained MorphDiff and IMPA. We also sampled 10 times for each model to ascertain the significance of the results. Figure 4 c illustrates that MorphDiff can effectively detect significantly changed CellProfiler features and outperforms IMPA for 7 out of 10 targets with the p-values less than 0.05 based on unpaired Welch t-test. In addition, we also validated whether our model can discriminate perturbations at the target level. We used Wasserstein distance to measure the diversity between DeepProfiler embeddings of perturbations at the target level. After computing the pairwise Wasserstein distance on target-level DeepProfiler embeddings, we found that the cell morphology generated by MorphDiff is highly consistent with ground truth as Figure 4 d shows. These results demonstrate that MorphDiff has good generalization ability and can learn more complex and in-depth relationships between drug perturbations and morphology. Moreover, MorphDiff can predict morphological changes and capture morphological diversity at the target level.

To further explore the potential utility of MorphDiff as a valuable tool in advancing drug discovery, we then focused on an important direction in drug discovery: identifying the MOA of a compound. We investigated whether cell morphology provides necessary information in the identification of drug MOA and whether the cell morphology inferred by MorphDiff also contains such information. Concretely, as demonstrated in Figure 4 e, we designed a framework for MOA retrieval and retrieval with cell morphology images. We constructed a database consisting of existing cell DeepProfiler embeddings treated by drugs with known MOAs. Given query cell morphology images, we also extracted the cell DeepProfiler embeddings and retrieved possible MOA based on the minimum distance between query morphology images and known morphology images corresponding to certain MOA. Due to the highly diverse and noisy properties of cell morphology, we adopted W-distance for DeepProfiler embedding retrieval to better account for the variance of cell morphology. The same framework can also be applied to MOA retrieval with gene expression by simply replacing the DeepProfiler embeddings with gene expression vectors and using the euclidean distance metric.

The retrieval experiments were conducted on the Target dataset. For the 68 drugs that act on 10 targets, we used different random seeds to divide them into two groups: one group of 46 (retrieval set) and another of 22 (query set). These 68 drugs correspond to 41 distinct Mechanisms of Action (MOAs), with some drugs associated with multiple MOAs. This complexity makes the retrieval task challenging. By calculating the W-distance (DeepProfiler embeddings) and MSE (Gene Expression) between the query set and retrieval set, we selected the top *k* closest drugs in the retrieval set as retrieval results for each query drug. If the correct MOA(s) is the same as or intersects with the MOA(s) of the top *k* nearest drugs, we regard it as a correct retrieval. We conducted top 5 retrieval of MOA(s) on ground truth DeepProfiler embeddings, MorphDiff-generated DeepProfiler embeddings, IMPA-generated DeepProfiler embeddings, gene expression and random retrieval results. To assess the significance of the experimental results, we randomly split the dataset 10 times with different random seeds. Figure 4 f demonstrates that MorphDiff not only outperforms gene expression but also significantly eclipses IMPA regarding retrieval accuracy, as indicated by the p-values of less than 0.05 and 0.01 respectively based on unpaired Welch’s t-test. The average accuracy of Morphdiff-generated output outperforms IMPA and gene expression-based retrieval by 29.1% and 9.7% respectively.

Furthermore, drugs with similar structures usually have the same target and MOA(s) in most cases [29]. However, it would be more interesting to investigate the drugs annotated with same MOA(s) but are structurally dissimilar or the drugs annotated with different MOA(s) but are structurally similar [30; 31] to improve biological safety and drug development flexibility. Therefore, we explored whether other modalities, such as gene expression and cell morphology, can provide complementary information in identifying these drugs, and we also investigated whether the cell morphology generated by MorphDiff on specific perturbations contains such complementary information.

For example, as shown in Appendix Figure 11, for drug embeddings, the closest drug embedding to *Clobetasol Propionate* is *clocortolone pivalate*, which has a different MOA *Steroid*. Besides, we can see that the drug embedding of *mometasone furoate* annotated with MOA *Glucocorticoid Receptor Agonist* is far away from other drug embeddings belonging to MOA *Glucocorticoid Receptor Agonist*. This indicates that the structures of *Clobetasol Propionate* resembles to *clocortolone pivalate* and differs from *mometasone furoate*. The structures of these drugs are shown in Appendix Figure 12. Meanwhile, as shown in Figure 4 g and Appendix Figure 11, even though ground truth DeepProfiler embeddings, gene expression as well as generated DeepProfiler embeddings exhibit seemingly different patterns, the closest drugs to *Clobetasol Propionate* for these different embeddings are all annotated with MOA *Glucocorticoid Receptor Agonist*. Furthermore, MorphDiff-generated and ground truth DeepProfiler embeddings both exhibit diversity between drugs from MOA *Glucocorticoid Receptor Agonist* and MOA *Steroid*, which is not captured by drug embeddings. At the same time, MorphDiff-generated DeepProfiler embeddings demonstrate higher consistency with ground truth DeepProfiler embeddings compared with IMPA as demonstrated in Appendix Figure 15 and Appendix Figure 16. In another case, as shown in Appendix Figure 13, for the drug *dichlorphenamide* annotated with MOA *Carbonic anhydrase inhibitor*, the corresponding closest drug in gene expression space is annotated with *Carbonic anhydrase inhibitor*, though their structures differ from each other. On the other hand, the closest drug to *dichlorphenamide* in ground truth and MorphDiff-generated morphological space is *fraxidin methyl ether* annotated with the same MOA. Interestingly, we can see that the structures of *fraxidin methyl ether* and *dichlorphenamide* are not similar as shown in Appendix Figure 14, which demonstrates that MorphDiff is potentially helpful in cases where researchers aim to discover an alternative drug structurally dissimilar with a known drug but of the same effect. Upon these observations, we found that the relationship between MOAs and gene expression, drug structure, and morphology is complex. None of these modalities can be used alone to perfectly predict the MOA thus the retrieval or prediction of MOAs is still a challenging task [10]. Integrating the shared and complementary information across these modalities will be a key to improving the identification of drugs’ MOA [8] and further advancing drug discovery. In summary, we validated the potential capability of MorphDiff in retrieving and identifying MOA of specific drug perturbation, with consistently better accuracy than IMPA and gene expression and comparable performance with ground truth. Furthermore, we showed that cell morphology contains complementary information in discriminating structurally similar drugs with different MOA and identifying structurally dissimilar drugs with the same MOA, with potential application in advancing phenotypic drug discovery.

## 3 Discussion

The exploration of cellular state transformation under genetic and drug perturbations enables numerous applications in drug discovery and biological research. Morphological profiling can cost-effectively capture thousands of features across perturbations by disease, mutation, or drug treatments and provide a unique view of cell state. Therefore, we propose MorphDiff, a diffusion-based generative model that can generate high-quality perturbed cell morphology conditioned with perturbed gene expression, efficiently leveraging the shared information between gene expression and cell morphology. We compared MorphDiff with several baseline methods and demonstrated its superiority with extensive experiments on two large-scale datasets. For genetic perturbation data, we analyzed the correlation between genes and morphology features, and the results show that MorphDiff effectively captures the pattern between gene expression and morphological profiling, establishing a bridge for studying the shared information between them. In the case of drug perturbations, MorphDiff successfully predicts the morphological changes caused by various drug perturbations. Furthermore, we applied the pre-trained MorphDiff model to unseen perturbations affecting specific targets and demonstrated its ability to capture diversity at the target level. This showcases the generalization capability of MorphDiff and its potential to facilitate the discovery of drug targets. Moreover, our experiments demonstrate the further application in drug discovery, enabling the investigation of drugs’ Mechanisms of Action (MOAs) through our advanced retrieval pipeline.

As for future work, first, MorphDiff is conditioned on perturbed gene expression, which means that MorphDiff cannot directly be applied to perturbations unmeasured by transcriptomic profiling experiments. Though L1000 assay [32] already profiles a vast number of diverse perturbations, there are still expectedly diverse new perturbations in the drug development process. Therefore, integrating MorphDiff with methods that can predict gene expression change under unseen perturbations, such as GEARS [33], is a possible extension. Second, the inference of MorphDiff is still costly as it is based on the diffusion model, which is known to be relatively slow in generating new samples. Therefore, the adaptation of more advanced sampling techniques for diffusion models in the future may further improve the efficiency of MorphDiff. Furthermore, incorporating more diverse input conditions, including but not limited to text description, drug structure, and chromatin accessibility besides gene expression, may further enhance the utility and generalizability of MorphDiff, making it a more powerful tool in predicting the cell morphology response under diverse perturbations. Lastly, MOA prediction remains ‘notoriously challenging’ as a major bottleneck in drug discovery, and no assay exists that can reliably be used to identify a majority of known MOA classes successfully [10]. Therefore, integrating multi-modal information, including but not limited to gene expression, morphological profile, and drug structures, may improve the MOA identification accuracy, and combining MorphDiff-generated cell morphology profile is a possible direction in achieving this goal.

The recent explosion of generative AI (GenAI) [11; 34; 35] has significantly transformed many areas. Vivid and high-resolution images and videos can now be generated with simple text descriptions at ultra speed and impressive quality. We anticipate that cell morphology, as a particular type of image, will also significantly benefit from the advancements of GenAI. Our work MorphDiff has demonstrated the potential of large-scale generative models in generating high-quality and high-fidelity cell morphology and explored several potential biological applications. It can be expected that large-scale pre-training on the increasingly available high-throughput cell morphology profiling datasets [5] with the rapidly developing GenAI method may continue to increase the precision and fidelity of cell morphology generation and further promote phenotypic drug discovery.

## 4 Method

### 4.1 Problem Definition

We collected *M* cell morphology 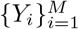 that correspond to *N* unique perturbations 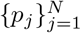 that can either be genetic perturbation or drug treatment. Specially, *p*_DMSO_ is the DMSO treatment that serves as control group with other perturbations and *Y*_DMSO_ is the cell morphology for the control group. The perturbation for cell morphology*Y*_*i*_ is denoted as *q*_*i*_ thus 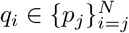. The L1000 gene expression corresponds to perturbation *q*_*i*_ is denoted as 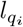. Our main objective is to train a model *f* that generates cell morphology conditioned on L1000 gene expression with high fidelity. Concretely, we aim to

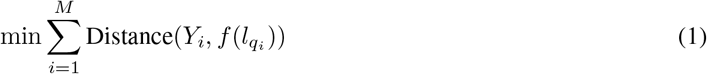

The cell morphology of the control group may serve as prior information for modeling the cell morphology after perturbation. Thus, *f* may also take *Y*_DMSO_ as input, which can be written as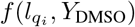.

### 4.2 MorphDiff Model

Following [11], MorphDiff is composed of two parts: Morphology VAE (MVAE) and Latent Diffusion Model (LDM). MVAE compresses the high-dimension multi-channel cell morphology images to low-dimensional latent representation with minimal information loss. LDM is trained to denoise from random Gaussian noise to lowdimensional latent representation by recursively adding Gaussian noise. In practice, a random vector is sampled from the Gaussian distribution and then passed to LDM while LDM sequentially denoises the random vector. High-quality predicted cell morphology can be decoded from the denoised output of LDM using the decoder of MVAE. Such approach offers a unique advantage: By compressing the high-dimensional cell morphology image to a low-dimension latent representation, MorphDiff is computationally efficient as sampling is performed on a low-dimensional space.

#### Morphology VAE (MVAE)

Morphology VAE (MVAE) comprises two parts, encoder *E* and decoder *D*. Given the cell morphology *Y* ∈ ℝ^*H*×*W*×5^, the encoder *E* encodes *Y* into a latent vector *z* = *E*(*Y*) and the decoder *D* reconstructs the cell morphology *Ŷ* = *D*(*z*). MVAE is trained in an adversarial manner following [36], that is, a patch-based discriminator *D*_*ϕ*_ is optimized to classify original cell morphology from the reconstructed morphology *D*(*z*). The patch-based discriminator is implemented with a multi-layer convolution neural network. It is a common practice in training VAE to regularize the latent *z* to be zero-centered and enforce small variance by introducing a regularizing loss term *L*_reg_. A Kullback-Leibler term *L*_reg_ between 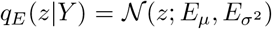 and standard normal distribution 𝒩 (*z*; 0, 1) is adopted to regularize the latent representation *z*. The overall objective of training MVAE can be summarized as:

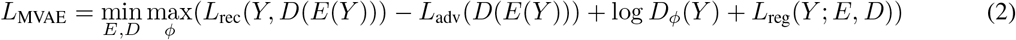

#### Learned Perceptual Image Patch Similarity (LPIPS) Loss in MVAE

The origin framework of VAE in the Stable Diffusion model [11] employs LPIPS loss [37] to minimize the difference between real images and generated images. The principle of LPIPS loss is to extract features of images with 5 conv layers from a pre-trained VGG network [38], and compute *l*_2_ distance between real features and generated features output from each layer. The integration of LPIPS will significantly improve the fidelity of VAE generation. However, as the pre-trained VGG model was trained with 3-channel images (RGB input), it is inconsistent with our 5-channel input (DNA, ER, RNA, AGP, Mito input). Therefore, We expand each channel to 3 dimensions to match the input shape of VGG. We compute LPIPS loss for each channel respectively and average across five channels. The formulation can be written as:

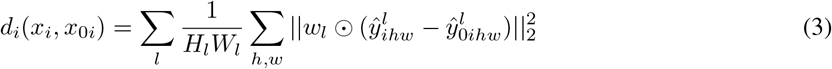

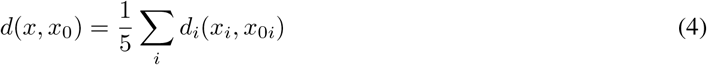

*l* represents *l*^*th*^ layer in 5-layers VGG and *w*^*l*^ is the parameters used to scale the activations. *x*_*i*_ and *x*_0*i*_ respresent the real image and generated image in *i*^*th*^ channel respectively, while 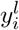 and 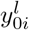 are adopted as corresponding features output from *l*^*th*^ layer. *H*_*l*_ and *W*_*l*_ are height and width of embeddings output from *l*^*th*^ layer.

#### Latent Diffusion Model (LDM)

Latent Diffusion Model (LDM) contains a noising and denoising process. Given a sample *z*_0_ ∈ *q*(*z*) where *q*(*z*) is the distribution of cell morphology latent representation. The noising process progressively adds Gaussian noise for *T* steps and obtains *z*_1_, *z*_2_, *…, z*_*T*_. The formulation of the noising process can be written as

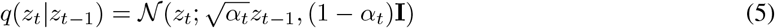

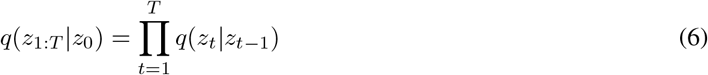

where *q*(*z*_*t*_|z_*t*−1_) can be interpreted as adding Gaussian noise to *z*_*t*−1_ parameterized by 1 −*α*_*t*_. We denote 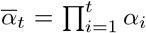. The denoising process aims to train a denoising network to recursively remove the noise from 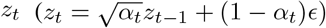 and *ϵ* ∼ 𝒩 (0, 1). Concretely, the denoising process can be formulated as follows:

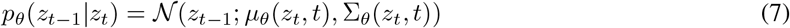

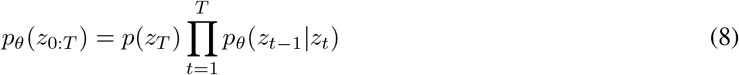

*µ*_*θ*_(*x*_*t*_, *t*) and Σ_*θ*_(*x*_*t*_, *t*) are the Gaussian mean and variance by the denoising network *θ*. Similar to other generative models, the loss function of LDM can be formulated as maximizing the log-likelihood of the samples generated belonging to the original data distribution (log(*p*_*θ*_(*z*_0_))). However, direct optimization is intractable as we need to integrate for *T* timesteps over the latent space. For L1000 gene expression *l*, we encode it as an intermediate representation and map it to UNet via cross-attention. Note that the encoding process is not learnable and it is a simple linear transformation, which is different from the training method in conditional latent diffusion [11]. This is because L1000 gene counts are already represented as vectors and processed in [8], so we believe it is better to map it into UNet directly. The encoder is represented as *EC*. Following DDPM [39], we can simplify the learning objective as:

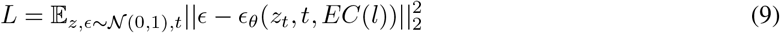

*z*_*t*_ is a noisy version of *z*_0_ at time *t* and *ϵ*_*θ*_(*t*)(*z*_*t*_, *t, l*) is a denoising network intended to predict *ϵ* from *z*_*t*_ conditioned on L1000 gene expression *l*. The cross-attention for *i*^*th*^ intermediate layer in UNet can be represented as 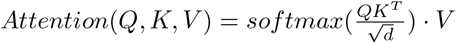, where 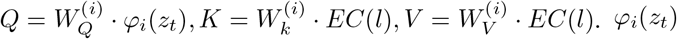 denotes a intermediate representation in UNet and 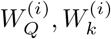 and 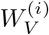 are learnable matrices. Following

DDPM [39], the detailed training and sampling algorithm of LDM is shown as follows:

##### Algorithm 1

Training Algorithm of MorphDiff

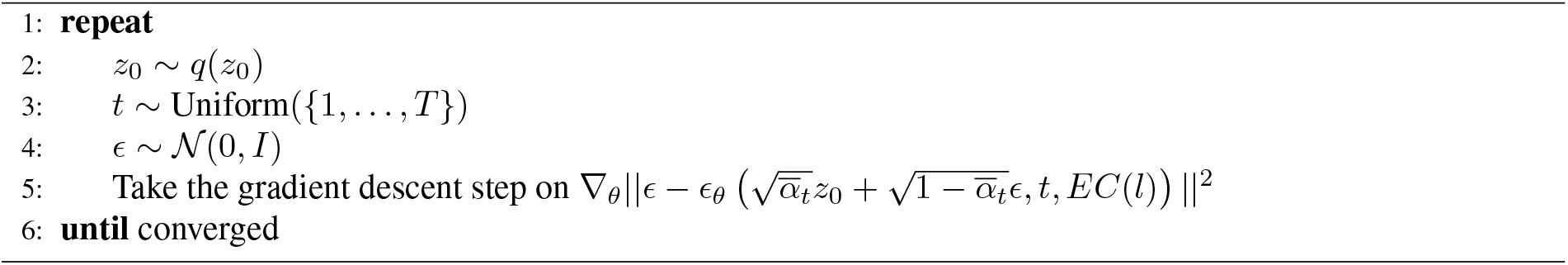

#### Implementation of LDM

LDM (*ϵ*_*θ*_(*z*_*t*_, *t, l*)) is implemented with the UNet [40] architecture augmented with an attention mechanism following [11]. Concretely, the base UNet architecture uses a stack of residual layers and downsampling convolutions, followed by a stack of residual layers with upsampling convolutions, with skip connections connecting the layers with the same spatial size. We use the UNet variant from [41] and replace the self-attention layer with alternating layers of (a) self-attention (b) position-wise MLP and (c) cross-attention layer. To inject the L1000 gene expression *l* as condition in the denoising UNet, we use an MLP to project *l* to an intermediate representation *τ*_*θ*_(*l*). The intermediate representation *τ*_*θ*_(*l*) is then mapped to each cross-attention layer 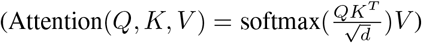 of UNet with

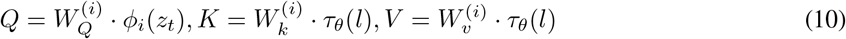

##### Algorithm 2

Buyer preferences for companies are influenced by factors extrinsic to the firm attributable to, and determined by, country-of-origin effects.

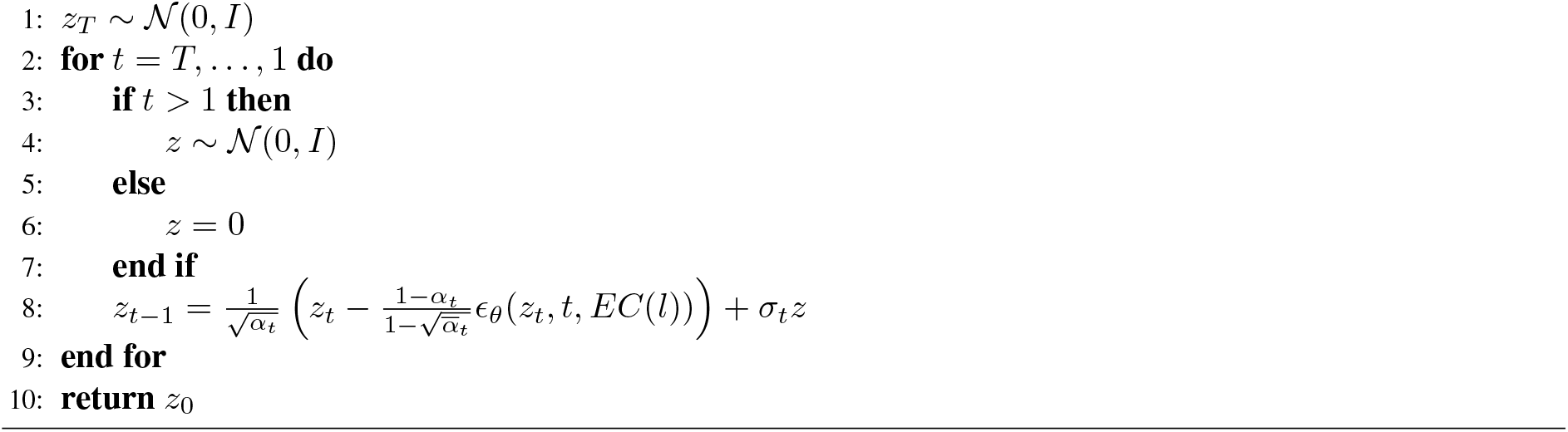

Sampling Algorithm of MorphDiff

*ϕ*_*i*_(*z*_*t*_) is the intermediate representation of the denoising UNet and 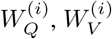 and 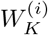 are learnable matrices.

### 4.3 Datasets

#### Drug treatment dataset

We collected the cell morphology images of cells treated with 960 different small molecules across 10 different plates (plate #25593, #25594, #25598, #25599, #26128, #26133, #26135, #26601, #26607, #26608) from [42], using the scripts from https://github.com/gigascience/paper-bray2017/blob/master/download_cil_images.sh. All the images consist of five channels, recording information from nucleus, Endoplasmic reticulum, Nucleoli and cytoplasmic RNA, Golgi and plasma membrane, and Mitochondria. The concrete correspondence between dye names, cell structures and CellProfiler names can be found in Appendix Table 1. We also collected bulk gene expression from https://broad.io/rosetta/ and L1000 gene expression in this dataset contain 977 genes. There are 961 categories totally, including 960 perturbations and control set.

#### Gene over-expression dataset

We collected the cell morphology images of cells treated with 130 different gene over-expression perturbations from the JUMP Cell Painting dataset [43] with the script from https://github.com/jump-cellpainting/datasets/. Each image consists of five channels, namely mitochondria (MitoTracker; Mito), nucleus (Hoechst; DNA), nucleoli and cytoplasmic RNA (SYTO 14; RNA), endoplasmic reticulum (concanavalin A; ER), Golgi and plasma membrane (wheat germ agglutinin (WGA); AGP) and the actin cytoskeleton (phalloidin; AGP). Meanwhile, we collected the bulk L1000 gene expression from L1000 database [32] with the same cell types and perturbations. L1000 gene expression in this dataset contain 12328 genes. The gene expression and the cell morphology images can thus be aligned according to the cell type and perturbation information. The total number categories is 131, including 130 perturbations and control set.

#### Image Processing

The images collected from [42] and [43] are all bulk-level cell plate images. To enable finegrained analysis at the single-cell image level. We used the CellProfiler software [19] version 4.2.5 to segment bulk-level cell plate images and get the single-cell level images. Concretely, we used the `IdentifyPrimaryObejects`and `IdentifySecondaryObjects`functions to identify individual objects(cells) in the images. The threshold strategy is set to global and the thresholding method is set to minimum cross-entropy.

#### Data alignment and split

We extracted the intersection between L1000 profiles and cell painting profiles at the perturbation level. The overall data processing pipeline is illustrated in Figure 1a. Both datasets are based on U2OS cells. The principle of selection is that all these perturbations can be simultaneously identified in both gene expression and morphology images, and there are a sufficient number of morphology images. Especially, the gene expression and morphology images are not paired and the number of morphology images are far more than gene expression. Because each perturbation normally only corresponds to 1 to 3 items of gene expression, we averaged gene expression for each perturbation and this averaged one are matched with all segemented single-cell images. After that, we split both datasets to three sets, which are training set, ID (in-distribution) set, and OOD (out-of-distribution) set. Firstly, we randomly picked 10 percent of perturbations for each datasets as OOD sets, resulting 13 perturbations for JUMP Cell and 96 perturbations for drug treatment dataset. The remained parts was split into training set (90 percent) and ID set(10 percent), which indicates that the training set and ID set contain both control and perturbation data.

#### Target dataset

We additionally selected 10 targets from [42], for which enough corresponding drugs and images could be collected for analysis and test. They are *NR3C1, EGFR, PPARG, CASP3, CDK2, CDK1, HMGCR, CA12, PRKACA* and *TOP2A*. Their corresponding perturbations are not present in our training set. After matching and cropping process mentioned above, we got 2222 images for *NR3C1*, 707 images for *EGFR*, 793 images for *PPARG*, 466 images for *CASP3*, 435 images for *CDK2*, 342 images for *CDK1*, 475 images for *HMGCR*, 327 images for *CA12*, 496 images for *PRKACA*. 51 images for *EGFR*. These datasets are unseen in the training set of Drug treatment dataset.

#### Morphological feature extraction

To evaluate the fidelity of the generated single cell morphology images, we used CellProfiler to extract features from the generated single cell morphology images, and compared them with the features from the real single cell morphology images. Some functions provided by CellProfiler are merely used for bulk-level image feature extraction, thus we selected several functions that can extract meaningful features of single-cell images. We employed `MeasureColocalization`(Measuring the correlation between any two channels), `MeasureGranularity`(Measuring the size and intensity of granular structures), `MeasureImageIntensity`(Measuring the intensity), `MeasureTexture`(Measuring the smoothness, coarseness, and regularity). Features with NaN values and zero variance will be removed. Meanwhile, features with duplicated meaning are also removed (e.g. Sum and Mean of intensity). After these processing and filtering steps, more than 200 CellProfiler features are kept for further analysis.

DeepProfiler [20] is a tool can extract dense embeddings from morphology images with the five-channel input. We used it following the handbook provided by authors. The handbook can be found at https://cytomining.github.io/DeepProfiler-handbook/docs/00-welcome.html.

#### Morphological feature selection based on discriminability of perturbations

CellProfiler Morphological features measure the cell morphology from multiple perspectives. Some morphological features may be highly correlated with the gene expression while some may not depend on gene expression or perturbation. Thus, to select most informative morphological features related to gene expression and perturbation for downstream analysis, we trained random forest classifier from morphological features to predict the corresponding perturbations. We set the number of estimators to 100 and selected the top 10 features with highest importance. The features with highest importance in the classifier are considered as the most discriminative of perturbations.

#### Morphological feature selection based on predictability

We selected most predictable morphological features to visualize the relationship between gene expression and cell morphological feature. Concretely, we computed the *R*^2^ score of each predicted cell morphological feature with respect to ground truth cell morphological feature. Only features with *R*^2^ score higher than 0.5 were kept. 36 features from the prediction of MorphDiff and 8 features from IMPA meet this criteria and 42 most predictable features are kept considering the overlap between two feature sets.

### 4.4 Baseline Methods

We used five common baselines in cell morphology generation and image translation to comprehensively benchmark the performance of our method. Unless otherwise stated, we used the default hyperparameters of these baseline methods.

#### MorphNet

MorphNet [14] is a computational approach that can infer images of a cell’s morphology from its gene expression. MorphNet contains two main components, the Variational Autoencoder (VAE) and Generative Adversarial Network (GAN). In the first training stage, the VAE is trained on the gene expression with ELBO loss to generate low dimensional gene expression embeddings. In the second training stage, the GAN network adapted from StyleGAN2 is used to generate cell images conditioned on the low dimensional gene expression embeddings. In the inference stage, the gene expression are first encoded with the VAE encoder and then the cell images are imputed through the GAN network.

#### IMPA

The Image Perturbation Autoencoder (IMPA) [3] is proposed to predict the cellular morphological effects of various kinds of perturbations using untreated cells as input. The model architecture is based on StarGANv2 [44]. Concretely, the model takes a control cell image *x*_*i*_ as input and the image is encoded into a dense representation. One perturbation embedding is sampled and concatenated with a random normally distributed vector. The concatenated embeddings are named as style embeddings and then fed into a perturbation encoder.

The encoded embedding will be used to condition every layer in the decoder. A perturbation discriminator will be employed to classify the specific perturbation from the decoder output. Meanwhile, a style encoder is trained to replicate the style vector from the decoder output image.

#### StarGANv1

StarGANv1 [18] performs conditional image-to-image translation across multiple domains using a single adversarial network. The objective is to learn a single generator for mapping across several different domains. During the training process, the discriminator is trained to minimize the classification error between real and generated data and maximize the classification accuracy on generated data across different domains. The generator aims to fool the discriminator with generated data on real/generated classification and maximize the classification accuracy across domains.

#### DRIT++

DRIT++ [17] is a powerful tool in domain adaptation and image translation. It encodes images onto a shared content space and domain-specific attribute space. The content encoders are trained to produce content embeddings not distinguishable by attribute discriminator. The cross-cycle consistency loss is adapted to exploit the disentangled content and attribute representations for cyclic reconstruction.

#### DMIT

DMIT [16] is a GAN framework developed for multi-condition image translation. The learning process of DMIT can be separated into disentanglement path and translation path. The input images will first be disentangled into the latent representation by an encoder-decoder architecture with conditional adversarial training. The generator will then learn multi-mappings across different domains by randomly performing cross-domain translation.

### 4.5 Evaluation Metrics

We denote the ground truth images and the generated images as **X** and **X**^′^ respectively and the number of images is *N*.

#### Freéhet Inception Distance (FID)

The Freéhet Inception Distance measures the similarity between two sets of images. It was proposed by [45] to quantify the similarity of generated images to real ones. Concretely, the generated images and real images are passed to the Inception V3 model pre-trained on ImageNet. The output encoding of real images and generated images are denoted as **E** and **E**^′^ with dimension (*N,* 2048) respectively (*N* is the number of images in the dataset and 2048 is the dimension of Inception V3 model encodings). The mean embedding vector of **E** and **E**^′^ are denoted as **m** and **m**^′^ and the covariance matrices **E** and **E**^′^ are denoted as **C** and **C**^′^. The final FID score is computed as 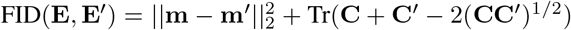. Tr denotes the trace of the matrix.

#### Inception Score

Inception score was introduced in [46] to measure the diversity and quality of the generated images. Mathematically, the inception score can be written as InceptionScore 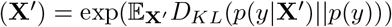. *y* is the marginal distribution of the Inception V3 model. First, the generated images are passed through the Inception V3 model to get the conditional distribution *p*(*y* | **x**), then, the marginal distribution *p*(*y*) is also computed by averaging *p*(*y* | **x**) over all the images. The KL divergence between *p*(*y*) and *p*(*y* | **x**) is calculated and averaged overall images. The exponential of the averaged KL divergence is used as the Inception score.

#### Density and Coverage

Density and coverage introduced by [3] provide additional measures for assessing the fidelity and diversity of the generated images. Basically, we assume that **E** = {**E**_1_, **E**_2_, *…,* **E**_*N*_} and 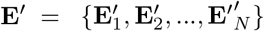 are the Inception V3 model encodings of the real images and generated images respectively. We can approximate a manifold of **E** as

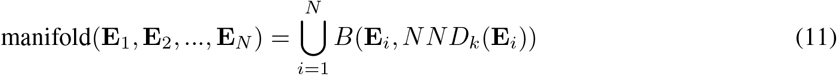

The sphere, denoted as **B**(*x, d*), is characterized by its center at the point *x* and its radius *d. NND*_*k*_(*x*) denotes the distance from point *x* to the *k*^*th*^ nearest neighbour *x* among excluding itself. Then, we can define density as

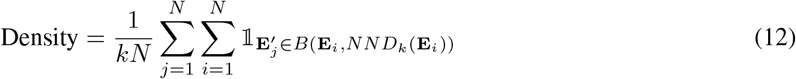

Intuitively, density quantifies the proportion of real neighborhoods containing generated samples. Meanwhile, coverage measures the extent that generated images recover the whole real image space. Coverage is defined as:

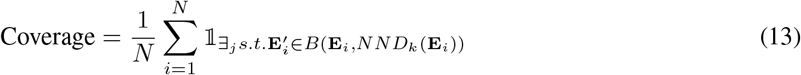

Intuitively, the coverage score will be higher if the generated images are close to the real images. We set *k* to 5 in our benchmark part.

#### Perturbation Classification

Perturbation classification is a straightforward way for measuring the specificity of the generated images with respect to certain perturbations. Concretely, for generated images belonging to the perturbation *A* and perturbation *B*. We label the images belonging to perturbation *A* as 1 and images belonging to the perturbation *B* as 0. We train a binary classifier using SVM with the image embedding extracted from DeepProfiler [20]. As some perturbations may not be distinguishable even on the ground truth images, we calculate a relative accuracy by dividing the accuracy of the generated images by the accuracy of the real images.

#### Coefficient of determination (*R*^2^ score)

The coefficient of determination (*R*^2^ score) [47] was created to evaluate prediction of various models and testing of hypotheses. It can assess how close the generated results are to the ground truth, based on the proportion of total variation of samples generated by models. Consider a ground truth set of n values marked *y*_1_, *y*_2_, *…, y*_*n*_, each associated with a predicted (or generated) value *g*_1_, *g*_2_, *…, g*_*n*_. Define residuals as *r*_*i*_ = *y*_*i*_ −*g*_*i*_ and 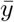 as the mean of ground truth set:

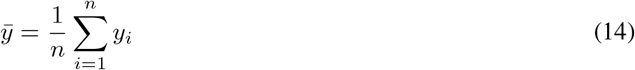

Then we can measure the variability of the data with two sums of squares formulas. The sum of square of residuals can be calculated as:

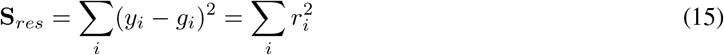

The total sum of squares is:

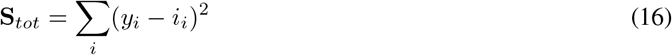

The general definition of *R*_2_ score is:

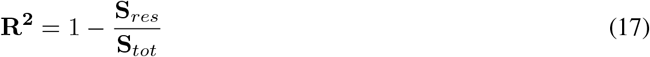

In the best case, the **S**_*res*_ = 0 and **R^2^** = 1. Generally, the *R*^2^ located in the interval (0, 1] means the degree of fitting and the closer the value is to 1, the better the performance of the model. Values of *R*^2^ being negative occur when the predicted data doesn’t fit the ground truth at all.

#### Wasserstein distance

Wasserstein distance [48] can measure the distance between different distribution. Intuitively, the origin of the Wasserstein distance lies in the optimal transport problem, which can be imagined as moving a pile of stones represented by probability distributions to form another target shape, while minimizing the cumulative distance of the movements. This is the central concern of the optimal transport problem. Let *µ* and *ν* be two distributions, and *d* is a way to calculate distance, then the Wasserstein distance can be defined as:

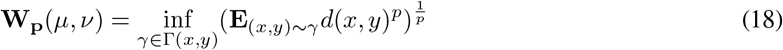

Where Γ(*x, y*) is the set of all couplings of *µ* and *ν*.

#### Pearson Correlation

Pearson correlation measures the linear relationship between two vectors. Concretely, given two vectors *X* and *Y*, pearson correlation *ρ* can be computed as

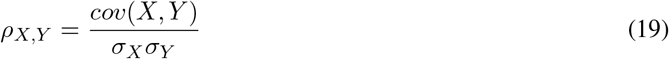

where *cov*(*X, Y*) is the covariance between *X* and *Y* and *σ*_*X*_ denotes the standard deviation of *X*.

#### F1 score

In this study, we adopt balanced F-score (F1 score) as the evaluation metric in Section 2.4. After chi-square test on CellProfiler feaures extracted from ground truth morphology, we term the features with p-value less than 0.05 as positive samples (regarded as significantly changed features), and term other features as negative samples. For generated morphology, we conduct same test on generated CellProfiler features, then we acquire predicted positive samples and negative samples generated by each method. In this way, we can calculate true positive (TP), false positive (FP) and false negative (FN) for each method. The F1 score can be calculated as:

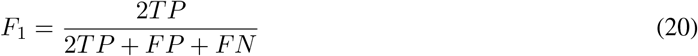

## 5 Data availability

All the data used in this work is publicly available. The Morphology images of drug perturbations dataset are from [42] and can be downloaded with the scripts from https://github.com/gigascience/paper-bray2017/blob/master/download_cil_images.sh. The corresponding L1000 gene expression data is collected from https://broad.io/rosetta/; The morphology images of genetic perturbations dataset are from the JUMP Cell Painting dataset [43] and can be downloaded with the script from https://github.com/jump-cellpainting/datasets/. The corresponding L1000 gene expression data is collected from [32].

## 6 Code availability

The reproducible data can be found in code ocean.

## 7 Acknowledgements

## 8 Author contributions

X.W., Y.F., Y.G., Y.L. and L.S. conceived the study. X.W., Y.F. designed the methodology. X.W., Y.F. and C.F. implemented the models. X.W., Y.F., C.F., K.L., K.D. and YX.L. ran the baseline experiments. X.W. and Y.F. interpreted the results. X.W., Y.F. performed analysis on the data. X.W., Y.F. and Y.G. organized and processed the publicly available data. X.W. and Y.F. visualized the experimental results. L.S., Y.L. supervised the research. X.W., Y.F. wrote the manuscript. Y.G. and Q.Y. help review the paper. All authors read and approved the final manuscript. L.S. and Y.L. founded the research.

