## Supplementary fig1-16 and table 1 for "Prediction of cellular morphology change under perturbations with transcriptome-guided diffusion model"

### Appendix Information for: Prediction of cellular morphology change under perturbations with transcriptome-guided diffusion model

Xuesong Wang*†[1,2], Yimin Fan*†[1,2], Yucheng Guo†[1], Chenghao Fu[2], Kinhei Lee[2], Khachatur Dallakyan[2], Yaxuan Li[2], Qijin Yin[1], Yu Li‡[2,4,5,6,7], and Le Song‡[1,3]

[1]BioMap Research, California, USA
[2]Department of Computer Science and Engineering, CUHK, Hong Kong SAR, China
[3]Mohamed bin Zayed University of Artificial Intelligence, Abu Dhabi, UAE
[4]The CUHK Shenzhen Research Institute, Hi-Tech Park, Nanshan, Shenzhen, 518057, China
[5]Institute for Medical Engineering and Science, Massachusetts Institute of Technology, Cambridge, MA, USA
[6]Wyss Institute for Biologically Inspired Engineering, Harvard University, Boston, MA, USA
[7]Broad Institute of MIT and Harvard, Cambridge, MA, USA

*Work was done while interning at BioMap.
†Equal first authorship.

8 Difference on morphological features between perturbed and control samples. . . . . . . . . . . . 10
9 MorphDiff predicts morphological changes on the target level. . . . . . . . . . . . . . . . . . . . . 11
10 (cont'd) MorphDiff predicts morphological changes on the target level. . . . . . . . . . . . . . . . . 12
11 Different modalities embeddings projected with UMAP. . . . . . . . . . . . . . . . . . . . . . . . 13
12 2D SMILES drug structure involved with Appendix Figure 11 . . . . . . . . . . . . . . . . . . . . 14
13 Different modalities embeddings projected with UMAP. . . . . . . . . . . . . . . . . . . . . . . . 15
14 2D SMILES drug structure involved with Appendix Figure 13 . . . . . . . . . . . . . . . . . . . . 16
15 The heatmaps describe the Wasserstein distance between pairwise morphological embeddings perturbed by various drugs. . . . . . . . . . . . . . . . . . . . . . . . . . . . . . . . . . . . . . . 17
16 The heatmap describes the Wasserstein distance between pairwise IMPA-generated morphological embeddings perturbed by various drugs. . . . . . . . . . . . . . . . . . . . . . . . . . . . . . . 18

### A Appendix Table

| Dye | Organelle or cellular component | CellProfiler |
|---|---|---|
| Hoechst 33342 | Nucleus | DNA |
| Concanavalin A/Alexa Fluor 488 conjugate | Endoplasmic reticulum | ER |
| SYTO 14 green fluorescent nucleic acid stain | Nucleoli, cytoplasmic RNA | RNA |
| Phalloidin/Alexa Fluor 594 conjugate, wheat germ agglutinin (WGA)/Alexa Fluor 594 conjugate | F-actin cytoskeleton, Golgi, plasma membrane | AGP |
| MitoTracker Deep Red | Mitochondria | Mito |

Appendix Table 1: **The CellProfiler channel name refers to the name assigned by the CellProfiler to each channel.** This table describes the cellular component corresponding to the CellProfiler channel name and the dye used for imaging.

### B Appendix Figures

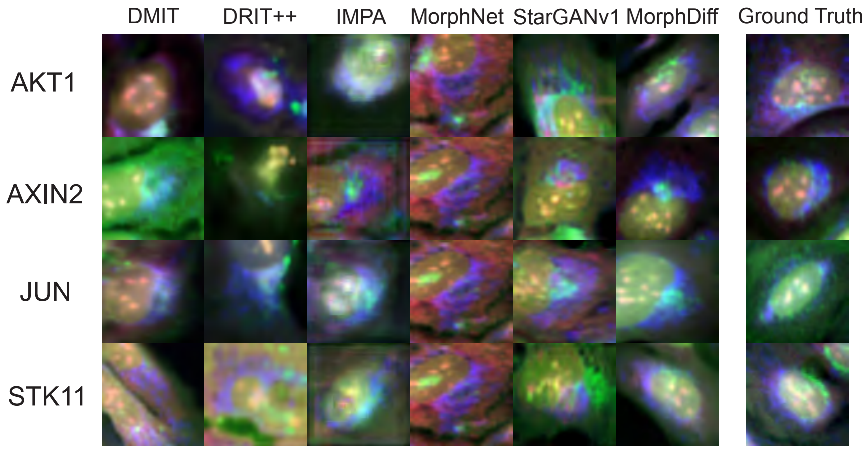

Appendix Figure 1: **Visualization of the generated samples of MorphDiff, ground truth, and several baselines on four genetic over-expression perturbations (AKT1, AXIN2, JUN, STK11).** We can observe that the generated samples of MorphNet are of relatively low quality and can not be distinguished across perturbations. The images generated by DRIT++ exhibit high variance across different perturbations, but the quality of the generated images is also low. StarGAN, DMIT and IMPA can generate relatively high-quality images, but the generated morphologies are unsatisfactory on some perturbations such as AXIN2 ang JUN. The visual quality of the images generated by the proposed method MorphDiff is the highest, and the generated images are the most visually similar to the ground truth images.

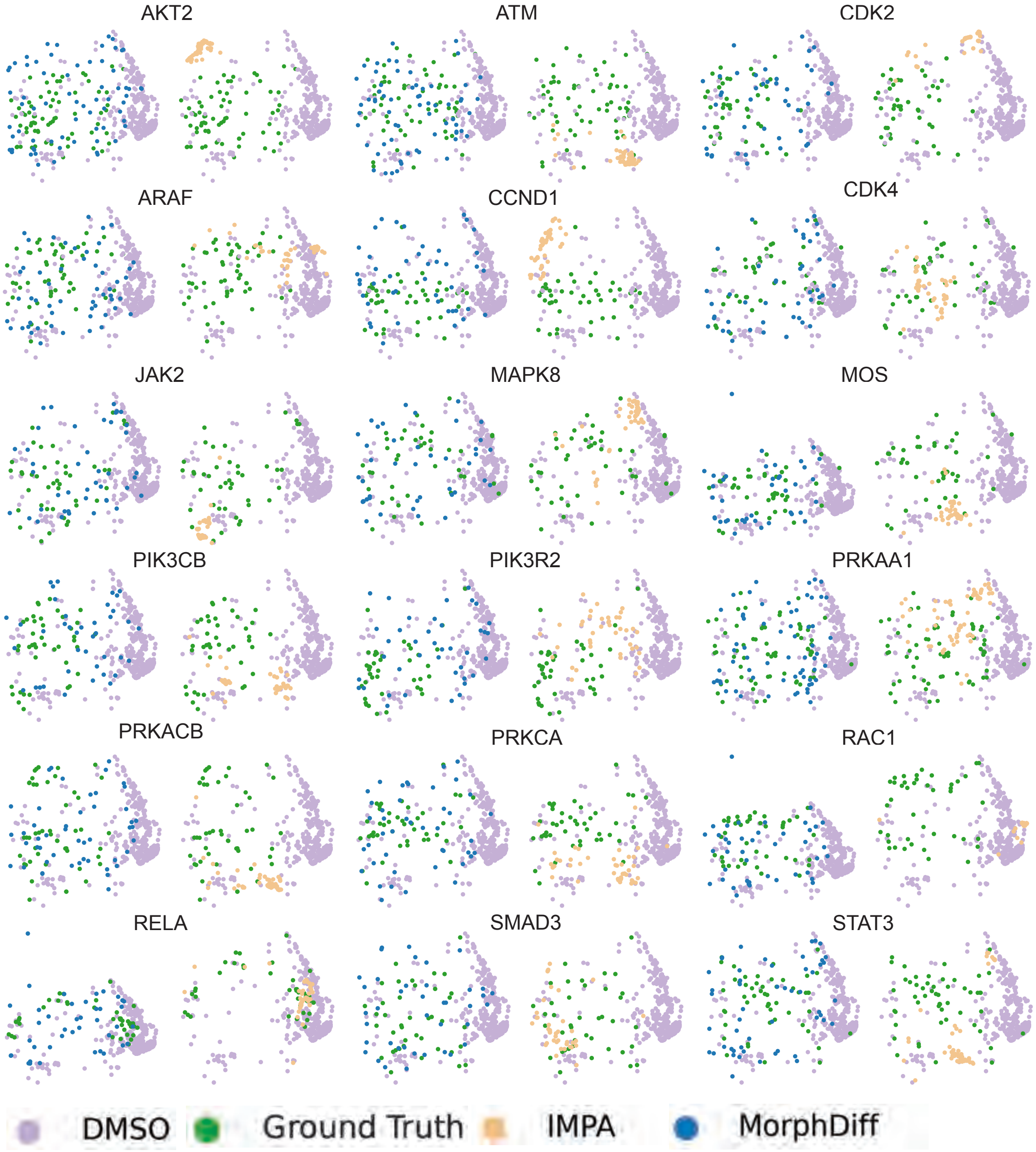

Appendix Figure 2: **UMAP visualization of the generated cell morphology CellProfiler feature on 18 genetic perturbations for IMPA and MorphDiff.** These perturbation morphology images are selected from the perturbations with the most images. This Appendix Figure corresponds to Figure 2 b in the main text.

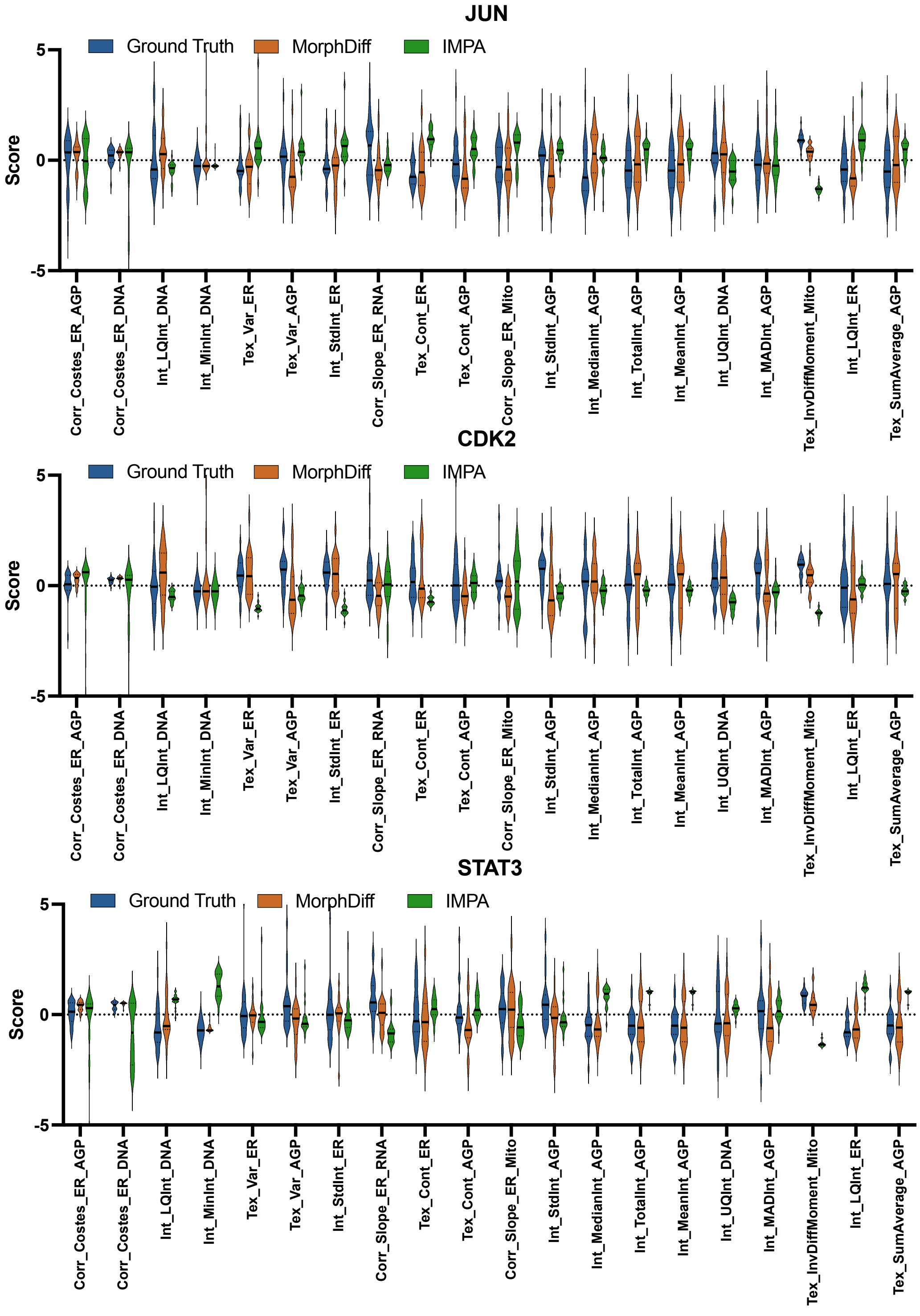

Appendix Figure 3: **Distribution of CellProfiler features between ground truth, IMPA-generated, and MorphDiff-generated morphology on JUN, CDK2 and STAT3 genetic perturbation.** *LQ* stands for *LowerQuartile*. *UQ* stands for *UpperQuartile*, *InvDiffMoment* stands for *InverseDifferenceMoment*, *Var* stands for *Variance*. *Tex* stands for *Texture*. *Corr* stands for *correlation*. *Int* stands for *Intensity*. This Appendix Figure corresponds to Figure 2 c in the main text.

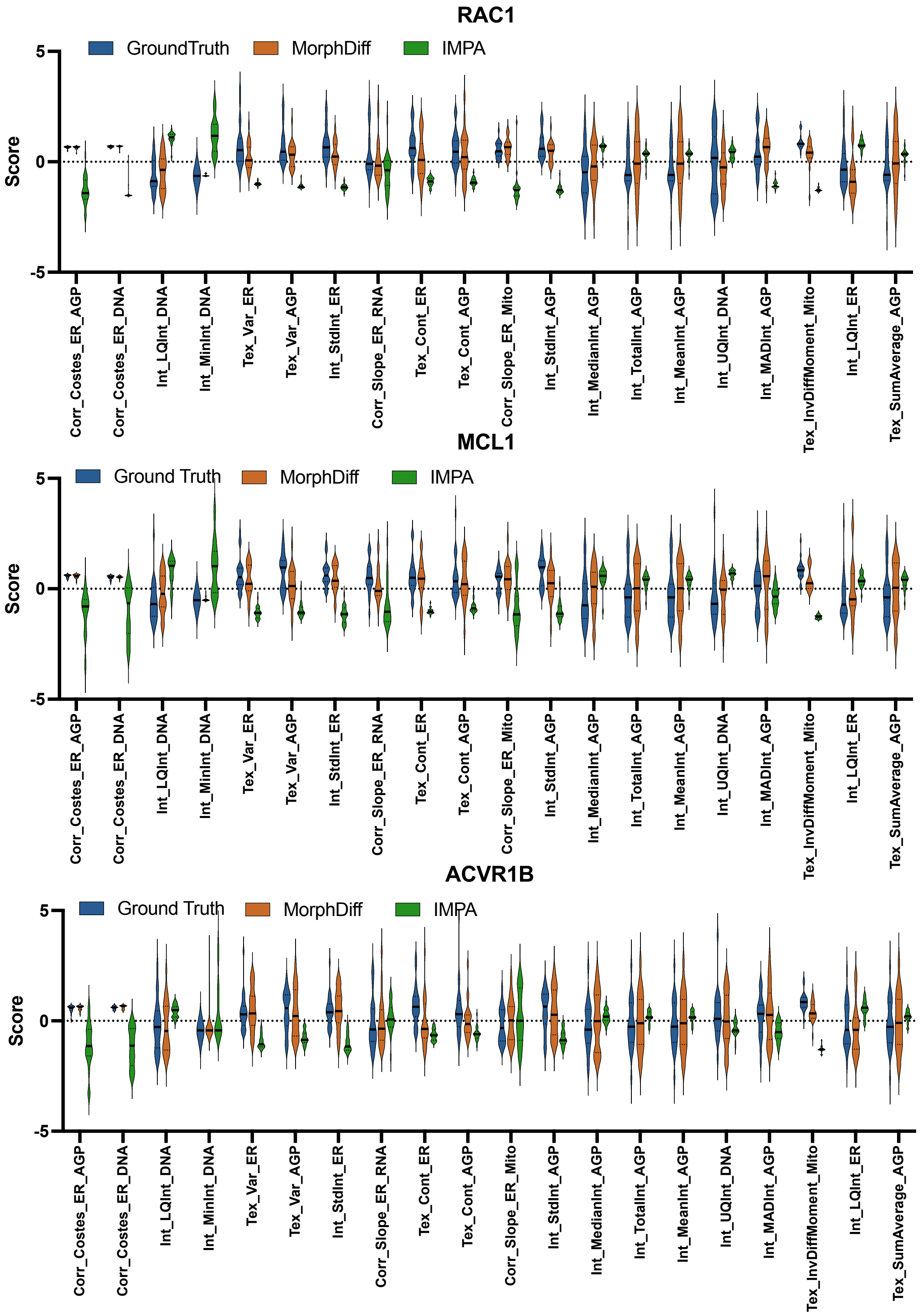

Appendix Figure 4: **(cont'd) Distribution of CellProfiler features between ground truth, IMPA-generated, and MorphDiff-generated morphology on RAC1, MCL1 and ACVR1B genetic perturbation.** *LQ* stands for *LowerQuartile*. *UQ* stands for *UpperQuartile*. *InvDiffMoment* stands for *InverseDifferenceMoment*. *Var* stands for *Variance*. *Tex* stands for *Texture*. *Corr* stands for *correlation*. *Int* stands for *Intensity*. This Appendix Figure corresponds to Figure 2 c in the main text.

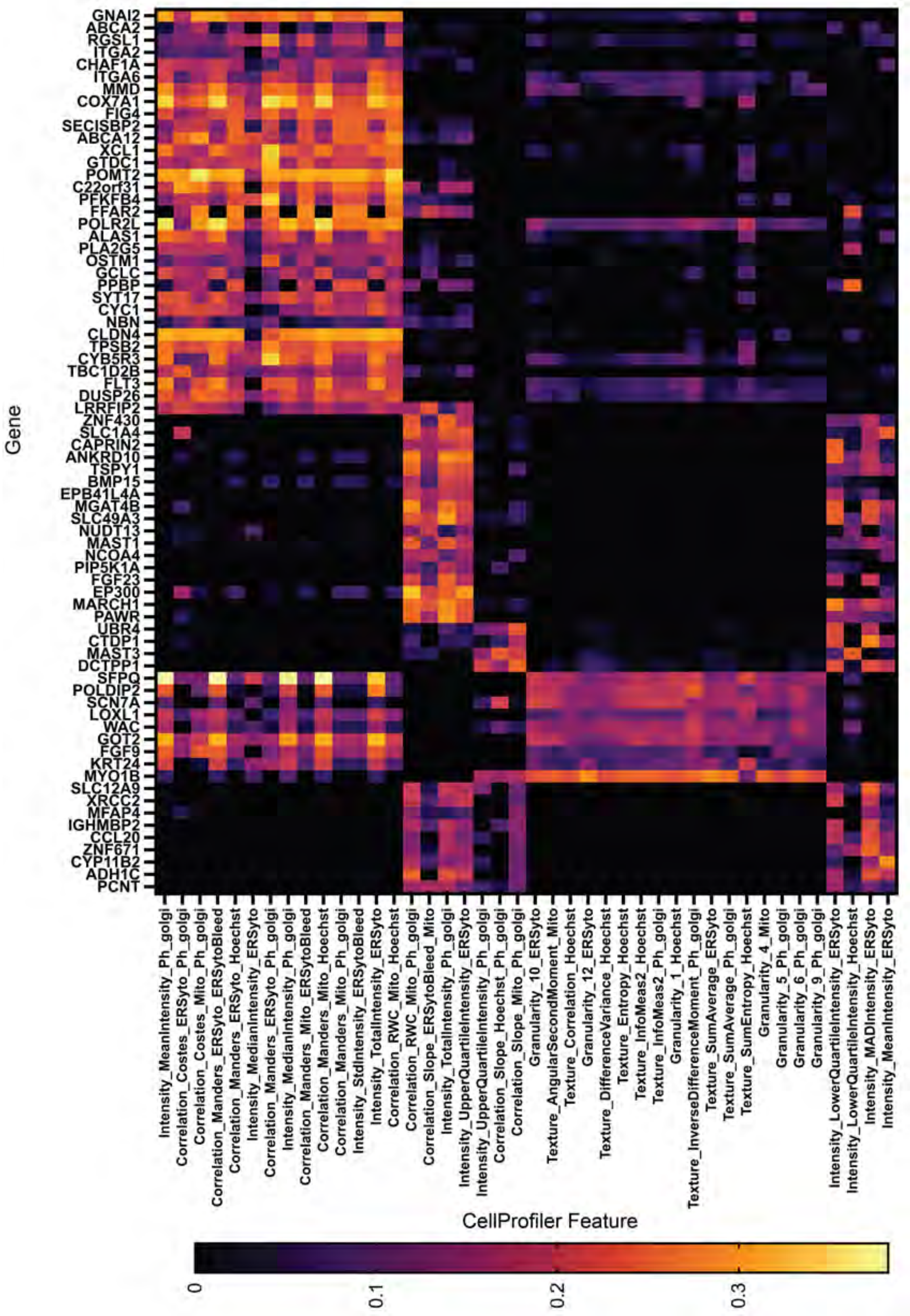

Appendix Figure 5: **Correlation heatmap between the gene expression and the CellProfiler Features from the ground truth cell morphology.**

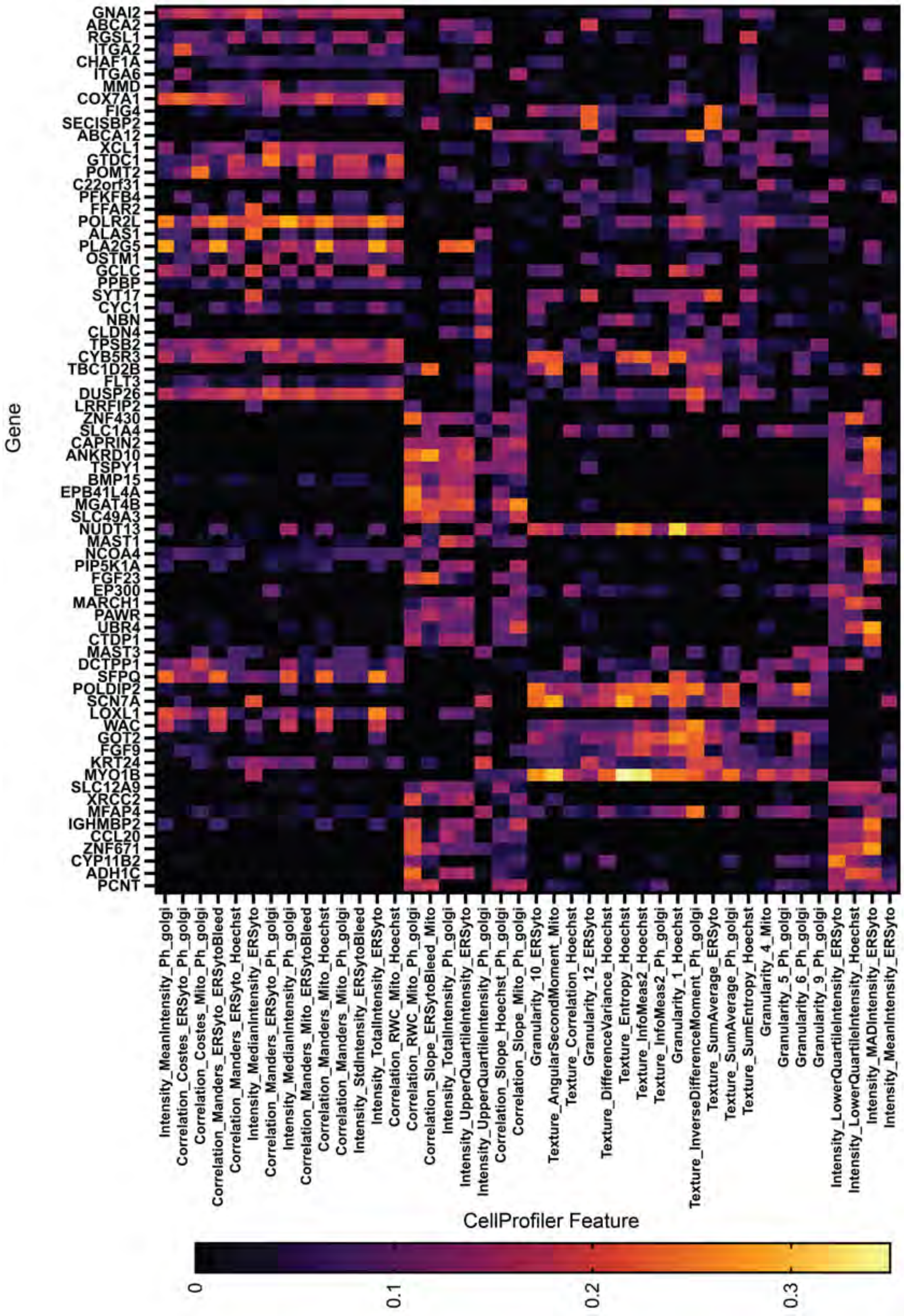

Appendix Figure 6: **Correlation heatmap between the gene expression and the CellProfiler Features from the predicted cell morphology from MorphDiff.**

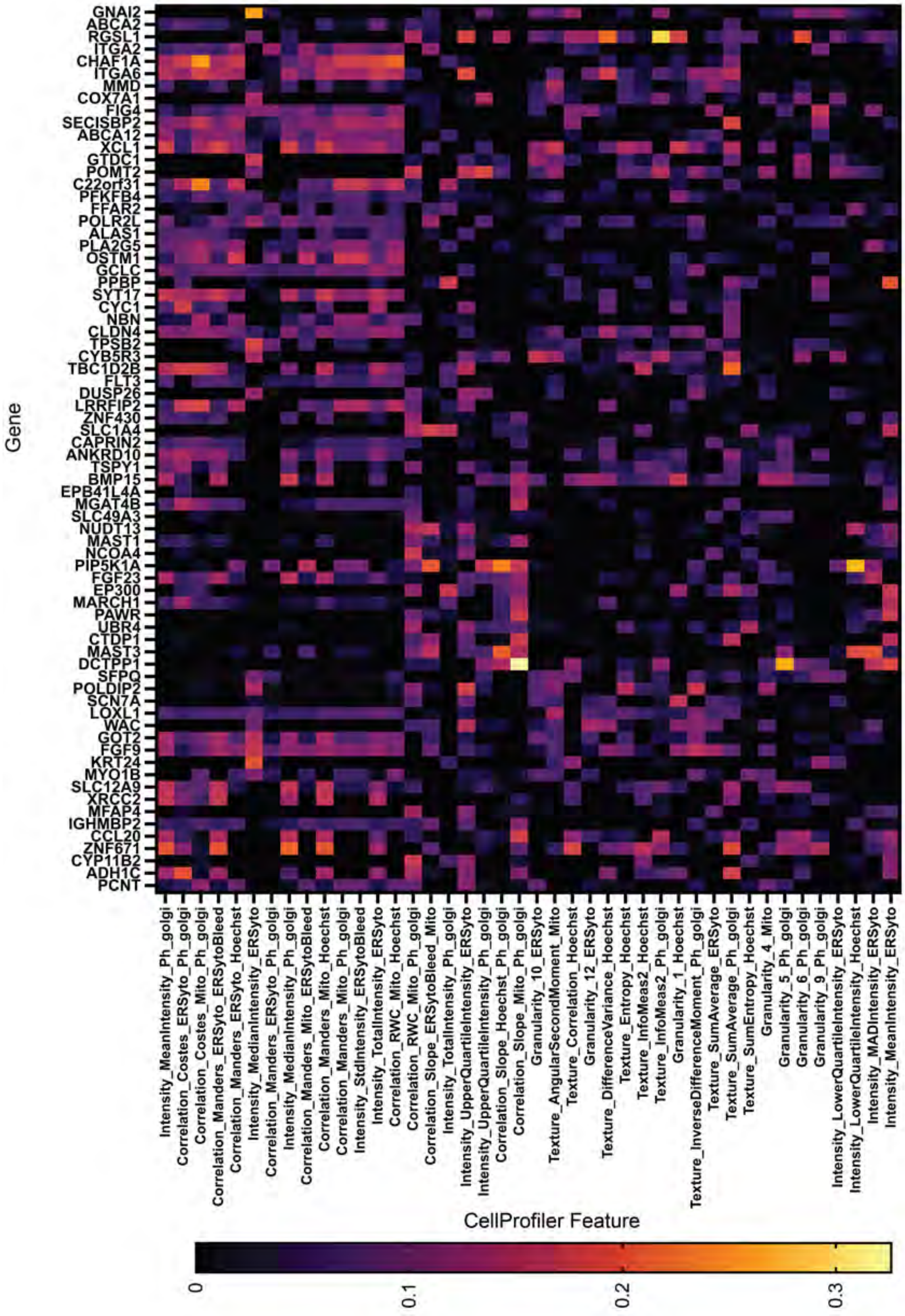

Appendix Figure 7: **Correlation heatmap between the gene expression and the CellProfiler Features from the predicted cell morphology from IMPA.**

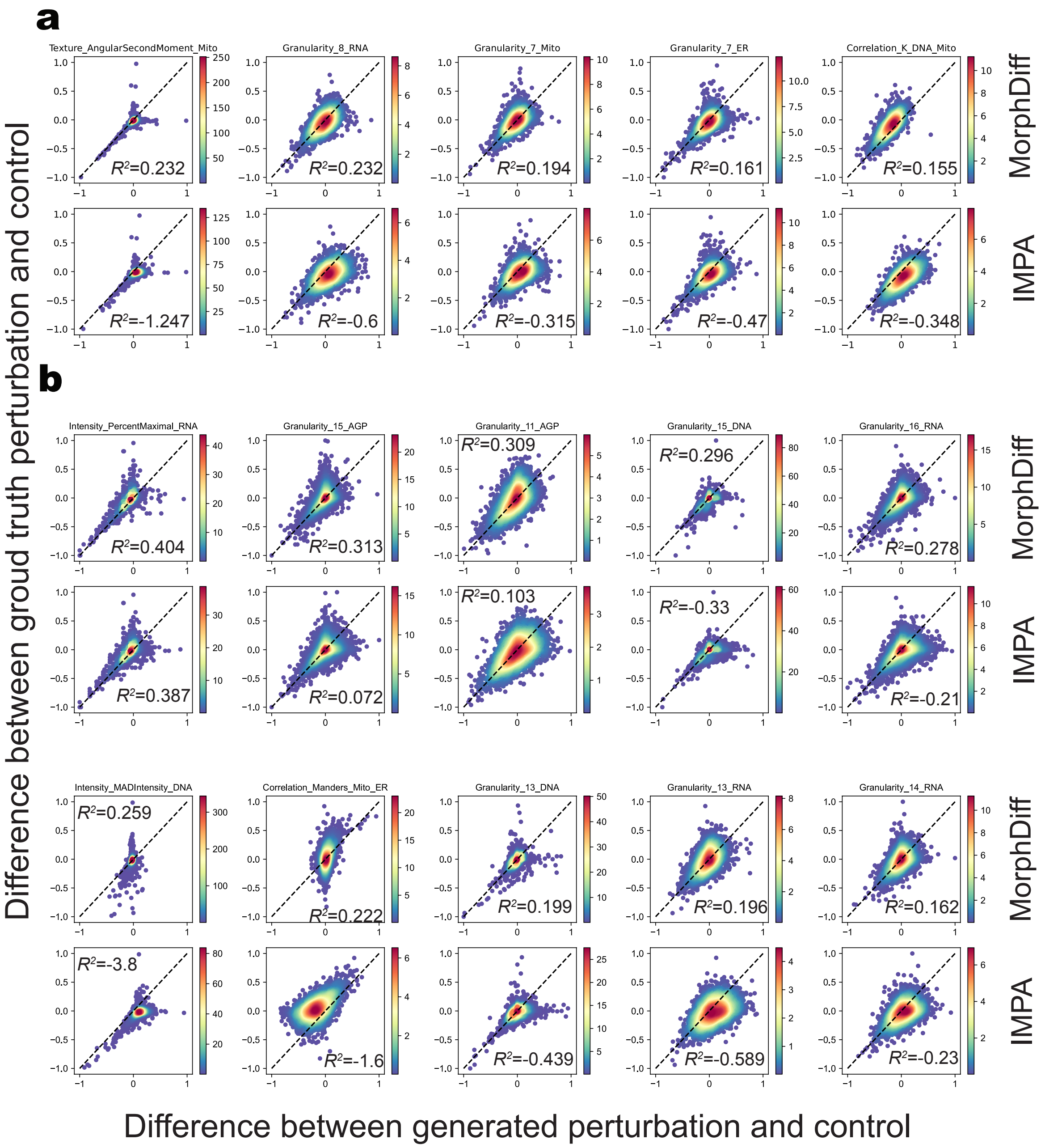

Appendix Figure 8: **Difference on morphological features between perturbed and control samples. a.** MorphDiff predicts cell changes on the Drug ID set. **b.** MorphDiff predicts changes in cells on the Drug OOD set. The x-axis displays the changes between the generated perturbation and the control, while the y-axis displays the changes in the ground truth perturbation and the control. The $R^2$ signifies the degree of fit between ground truth and generated changes.

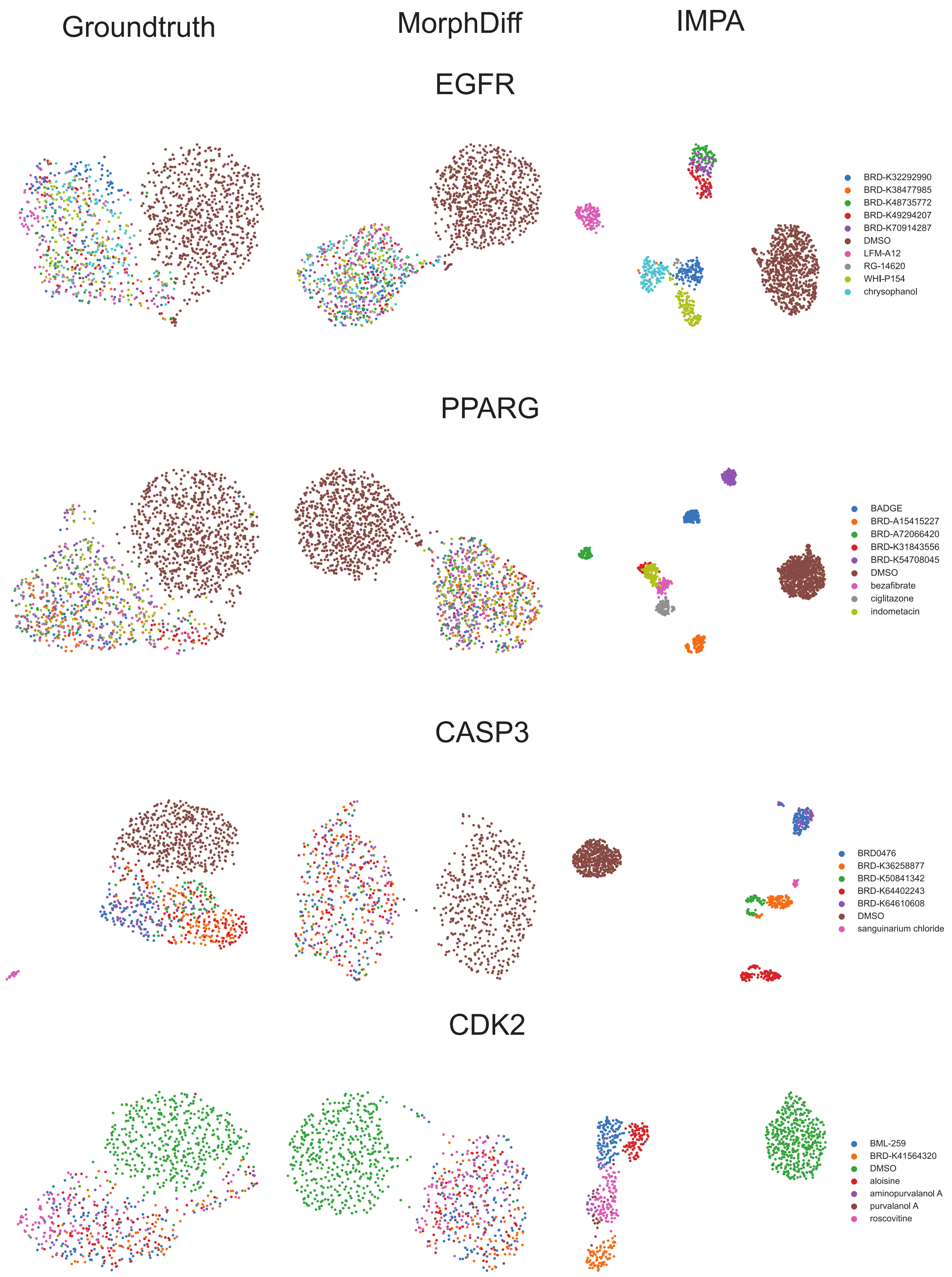

Appendix Figure 9: **MorphDiff predicts morphological changes on the target level.** Morphological features were extracted from both ground truth and generated images. The perturbed morphological features can be distinguished from the control, and the drugs targeting the same target cause similar changes in morphological space. The images generated by MorphDiff are consistent with this distinction, while IMPA generates different clusters consisting of different perturbations.

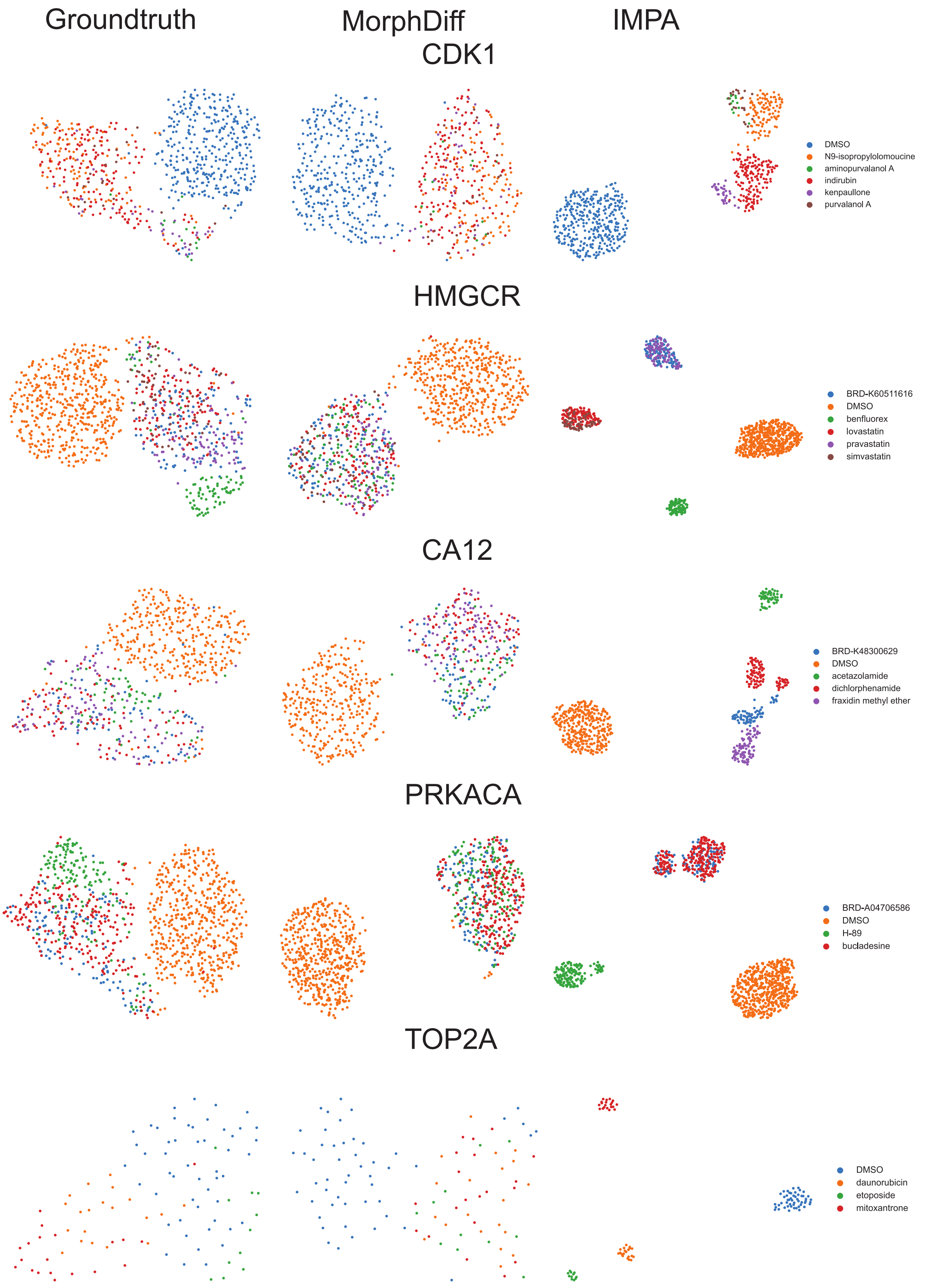

Appendix Figure 10: **(cont'd) MorphDiff predicts morphological changes on the target level.** CellProfiler features were extracted from both ground truth and generated images for the other five targets.

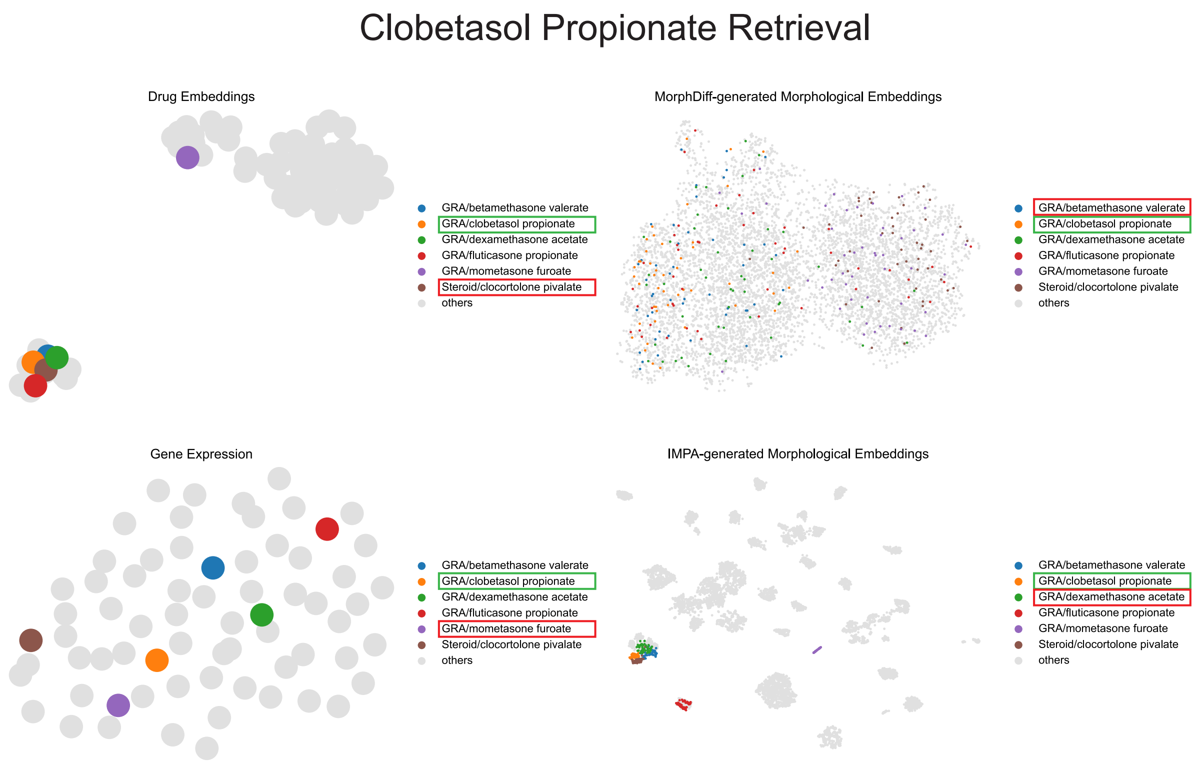

Appendix Figure 11: **Different modalities embeddings projected with UMAP.** The drug in the green box is *clobetasol propionate* annotated with GRA, and the drug in the red box is the closed drugs based on Wasserstein distance (DeepProfiler embeddings) or MSE (gene expression). GRA and Steroid are two types of MOAs. GRA is the abbreviation of *Glucocorticoid Receptor Agonist*.

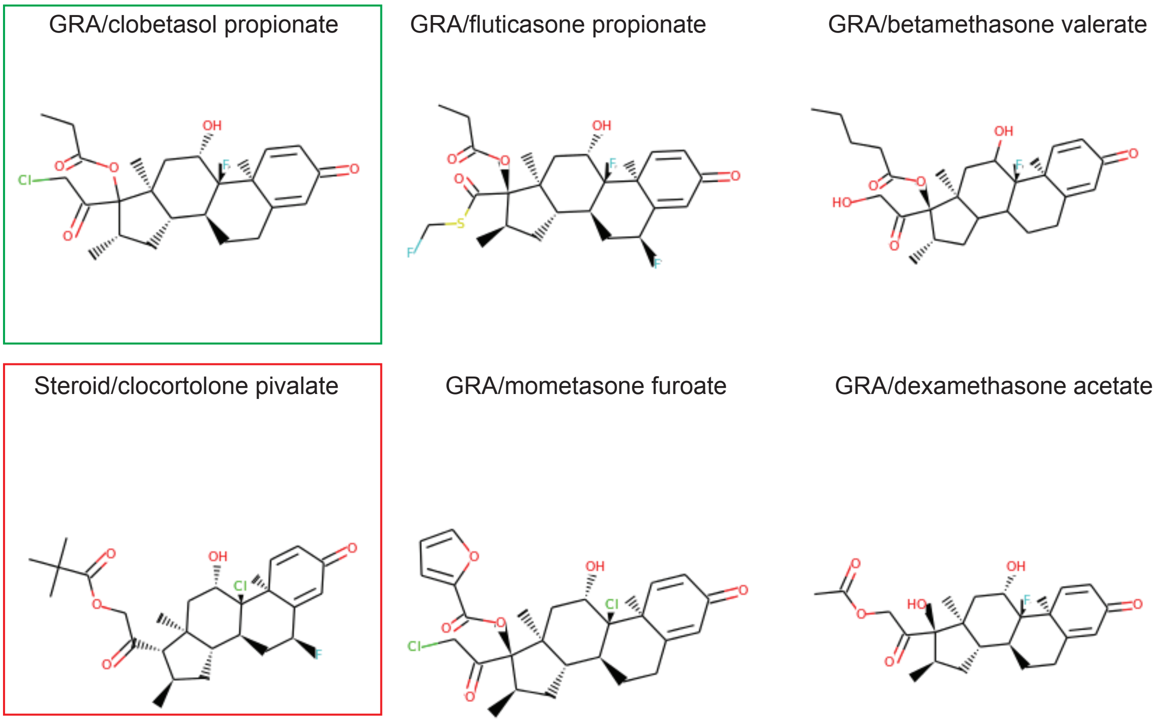

Appendix Figure 12: **2D SMILES drug structure involved with Appendix Figure 11, generated from `http://hulab.rxnfinder.org/smi2img/`.** The Structure in the green box is the query drug, and the Structure in the red box is the false retrieval result.

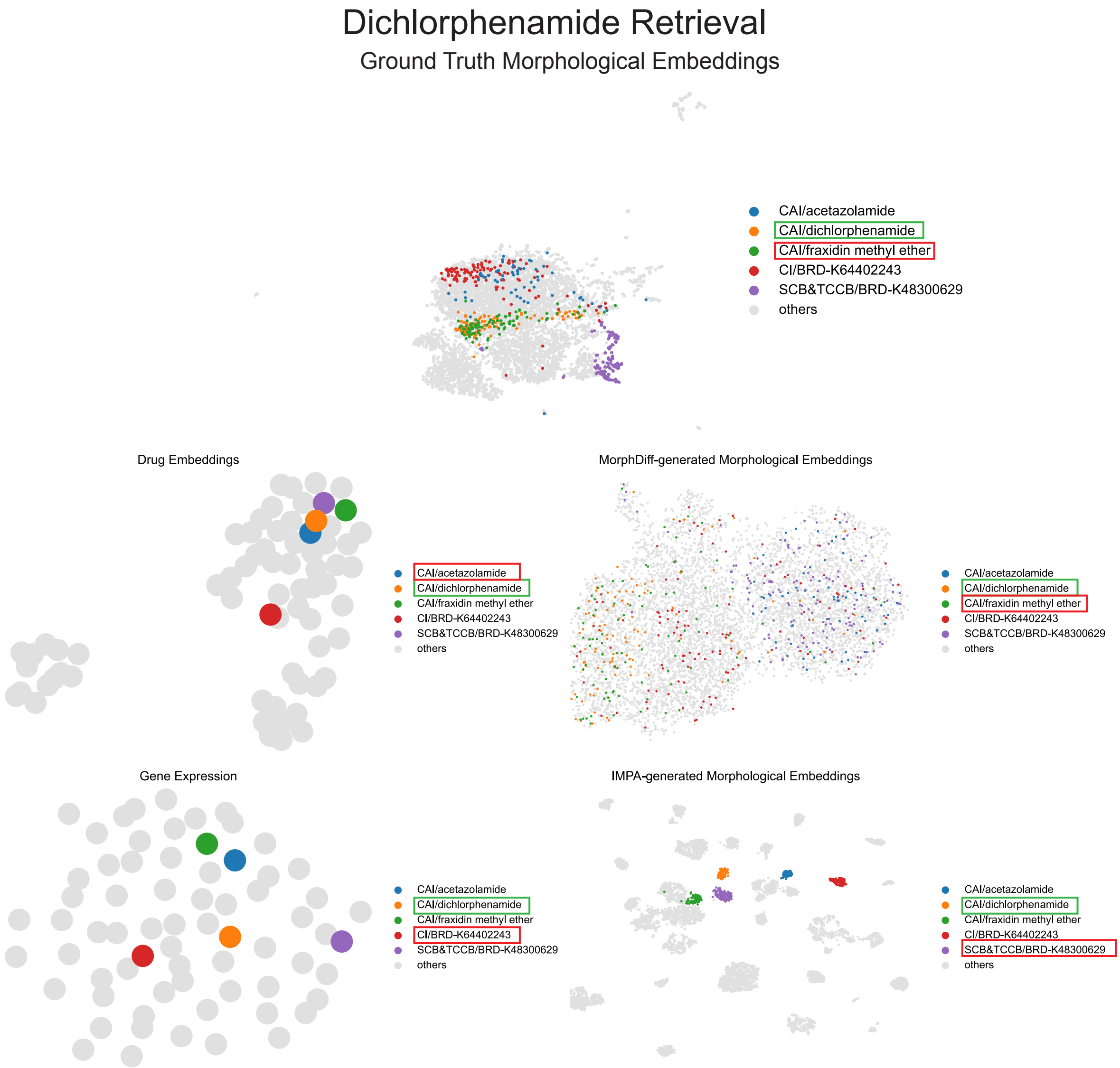

Appendix Figure 13: **Different modalities embeddings projected with UMAP.** The drug in the green box is *Dichlorphenamide* annotated with CAI, and the drug in the red box is the closed drugs based on Wasserstein distance (DeepProfiler embeddings) or MSE (gene expression). CAI is the abbreviation of *Carbonic Anhydrase Inhibitor*. CI is the abbreviation of *Caspase inhibitor*. SCB&TCCB is the abbreviation of *Sodium channel blocker & T-type calcium channel blocker*.

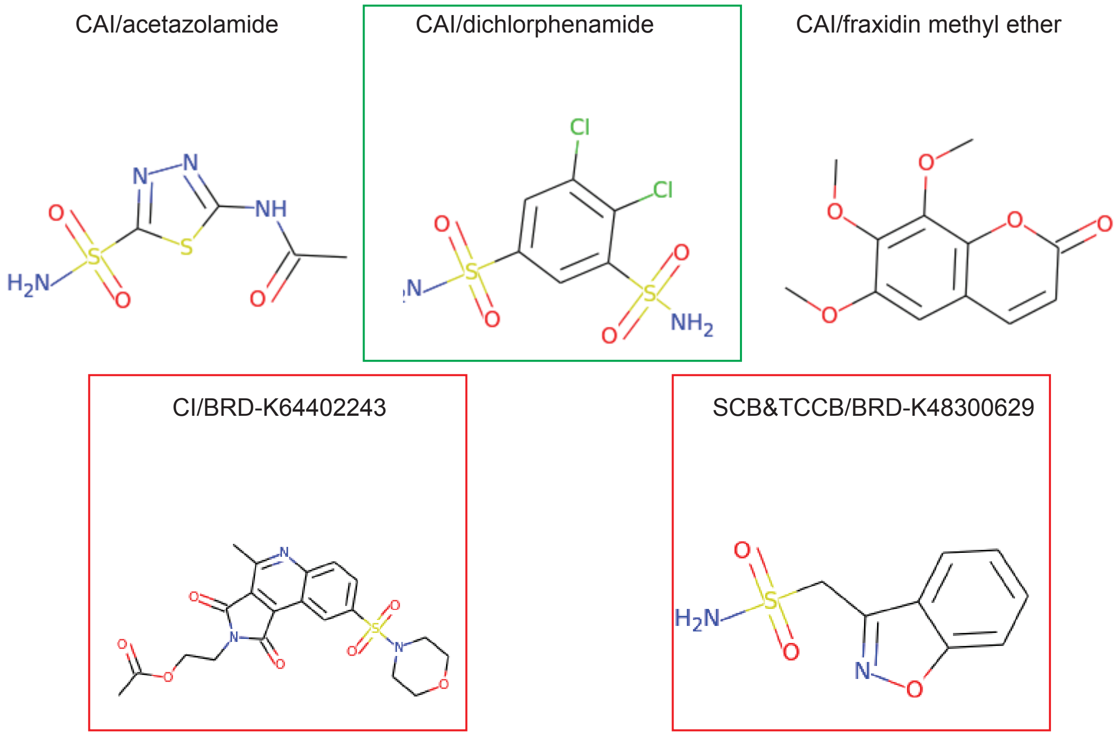

Appendix Figure 14: **2D SMILES drug structure involved with Appendix Figure 13, generated from `http://hulab.rxnfinder.org/smi2img/`.** The Structure in the green box is the query drug, and the Structure in the red box is the false retrieval result.

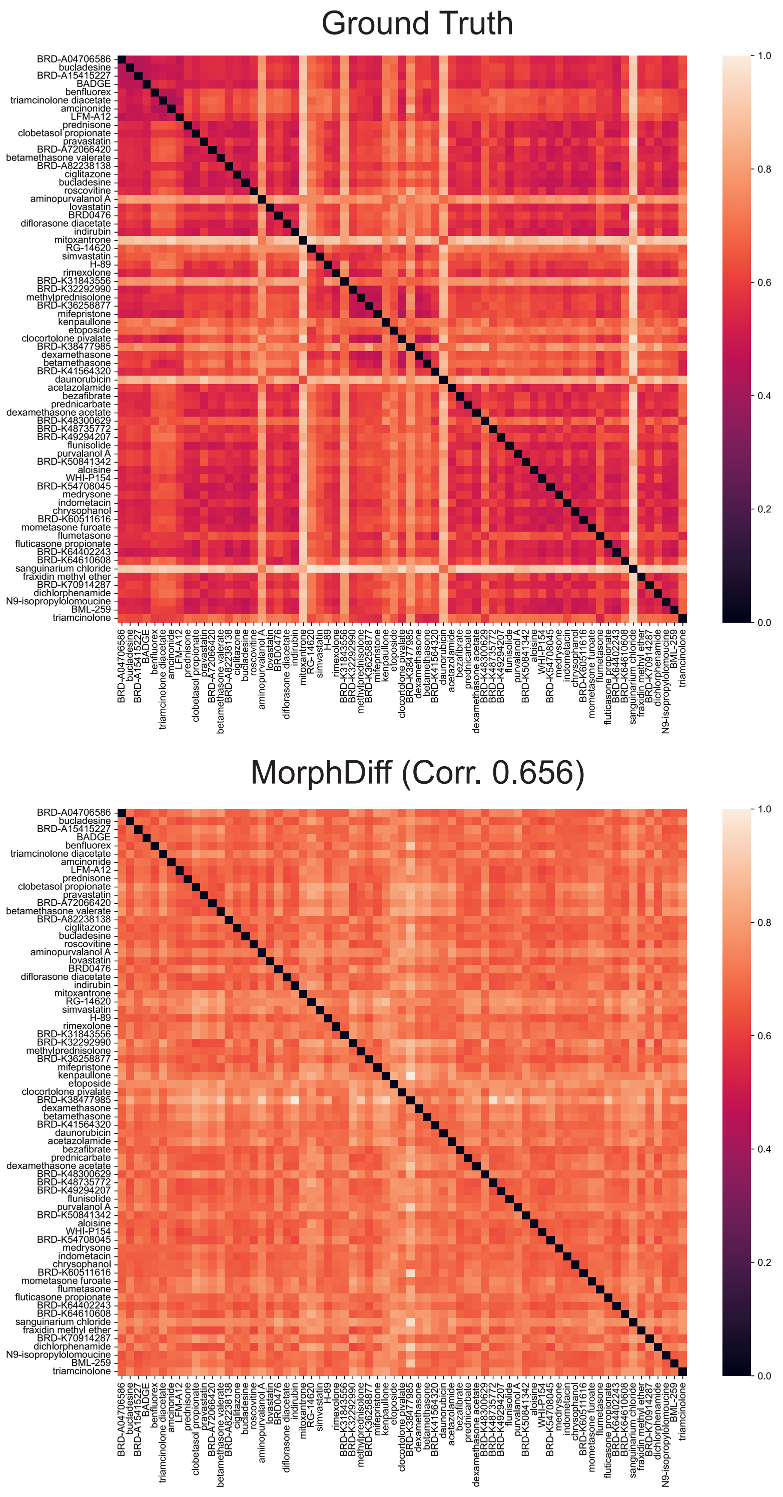

Appendix Figure 15: **The heatmaps describe the Wasserstein distance between pairwise morphological embeddings perturbed by various drugs.** The correlation between the heatmap calculated with ground truth and MorphDiff-generated morphological embeddings is 0.656.

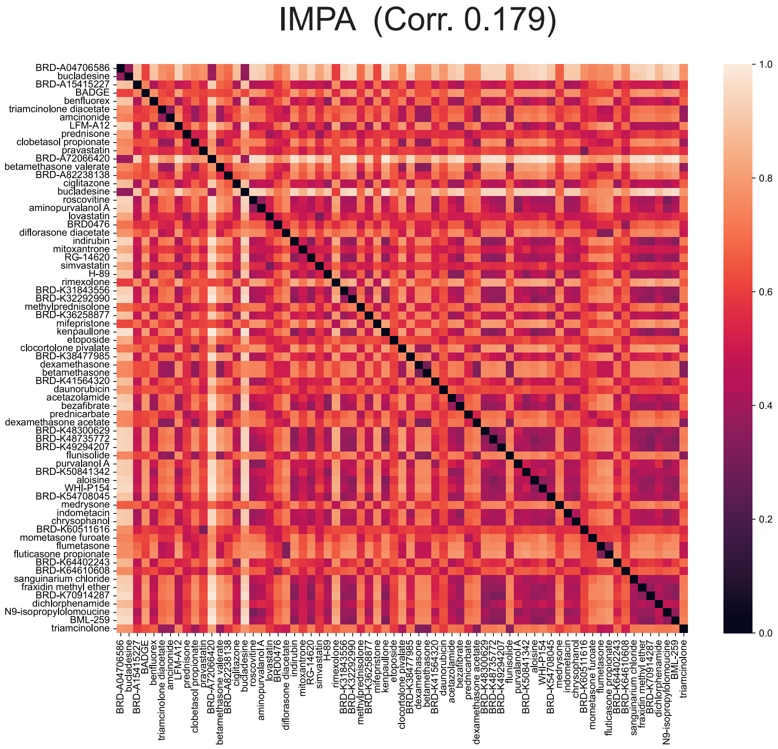

Appendix Figure 16: **The heatmap describes the Wasserstein distance between pairwise IMPA-generated morphological embeddings perturbed by various drugs.** The correlation between the heatmap calculated with ground truth and IMPA-generated morphological embeddings is 0.173.
